# Development and evaluation of a cost-effective, mid-density SNP array as a sorghum community genotyping resource

**DOI:** 10.64898/2026.02.20.706663

**Authors:** Vivek Kumar, Robert R Klein, Benjamin Kaufman, Noah D Winans, Daniel Crozier, William L Rooney, Melanie Harrison, Chad Hayes, Marcela K Tello-Ruiz, Nicholas Gladman, Christopher Olson, Gloria Burow, Sarah Sexton-Bowser, Somashekhar Punnuri, Joseph Knoll, Jeff Dahlberg, Doreen Ware

## Abstract

The development of accessible and cost-effective genotyping platforms is essential to accelerate genetic gain in crop improvement. To address the U.S. sorghum community’s need for a standardized, mid-density genotyping resource, we developed and validated a targeted single-nucleotide polymorphism (SNP) array using the PlexSeq™ next-generation sequencing (NGS) platform. The resulting genotyping array includes 2,421 SNPs spanning all ten Sorghum bicolor chromosomes and integrates trait-linked and quality control markers selected by public and private stakeholders. Genotyping 2,726 diverse accessions, including the Sorghum Association Panel (SAP), demonstrated high call rates (>90% for most samples and markers), low missing data, and accurate resolution of population structure consistent with prior whole-genome studies. In comparative genomic prediction analyses, the mid-density array performed equivalently to high-density genotype-by-sequencing (GBS) platforms for key traits such as grain yield and plant height across multi-environment trials. Designed for broad utility in breeding pipelines, the array enables marker-assisted selection, genomic prediction, identity verification, and germplasm quality control. Moreover, its adoption by the USDA National Plant Germplasm System facilitates the curation of genebanks and the management of core collections. This community-driven genotyping platform offers a scalable, reproducible, and customizable tool to support molecular breeding in sorghum and underscores the value of targeted marker systems in resource-optimized crop improvement programs.

## Introduction

Sorghum [*Sorghum bicolor* (L.) Moench], a member of the grass family, Poaceae, is the fifth most important cereal crop globally, following wheat, rice, maize, and barley. Cultivated across over 41 million hectares globally, with a production exceeding 62 million tons, sorghum is a cornerstone of food security, particularly in the arid and semi-arid regions of Africa and Asia (FAO 2021). The crop’s historical journey began approximately 6,000 years before present (BP) in present-day Sudan, where it formed a vital component of the African food complex that facilitated human expansion and migration (Mann et al. 1983). Through centuries of human-mediated selection, sorghum has diversified into thousands of distinct ecotypes, adapted to various climates and serving specialized uses, including gluten-free grain, livestock forage, industrial raw material, and a source for cellulosic and ethanol-based biofuels. Sorghum’s inherent tolerance to drought and heat stress, coupled with a versatile growth habit, positions sorghum as a critical resource for global food and agriculture security in the face of escalating climate change (Mwamahonje et al. 2024; Khalifa and Eltahir 2023).

To date, the United States is the largest producer of sorghum with over 4.7 million tons harvested on over 1.8 million hectares (FAO 2022) with production predominantly in the semi-arid regions of Kansas, Texas, Oklahoma, Nebraska, and South Dakota. While traditional markets include ethanol, livestock feed, and exports, a recent surge in demand for gluten-free and plant-based foods has expanded sorghum’s role in the consumer food industry (Tanwar et al. 2023; Whole Grains Council 2025).

Sorghum’s introduction to the U.S. is thought to have occurred in the 17th century, where it was initially cultivated as a crop for animal feed. In the late 19th and early 20th centuries, U.S. production was based on a limited number of cultivars such as Milo and Kafir, and the first critical advancement in grain yield was the development of shorter combine-height sorghum suitable for mechanized harvesting in the 1940s. The second critical advancement in grain yield was the discovery of cytoplasmic male sterility (CMS) and rapid adoption of sorghum hybrids in the 1950s (McCollough 1972; Duncan et al. 1991). In the early 1970s, the TAES-USDA Sorghum Conversion program was introduced as a joint venture between the Texas Agricultural Experiment Station (TAES) and the USDA Agricultural Research Service (USDA-ARS) to convert tropical sorghum lines into cultivars adapted for temperate zones. This innovative germplasm conversion program unlocked the vast amount of genetic diversity in tropical sorghum that was largely inaccessible to temperate-zone sorghum improvement programs and resulted in cultivars with enhanced traits such as disease and insect resistance, drought tolerance, and better grain quality and yield. Despite its importance, U.S. sorghum grain yields have plateaued in recent decades, a trend often attributed to comparatively lower research investment than other major cereals like maize and sorghum’s cultivation in marginal environments (Figure 1). To counter this trend within the US, sorghum grower-supported organizations, such as the National Sorghum Producers and the United Sorghum Checkoff Program, have played key roles in advancing the crop, with the former focused on policy advocacy and industry representation, and the latter investing in research, education and market development. Owing in part to these grassroot efforts, the top three markets in the US have been ethanol, livestock feed, and exports. However, there has been a recent surge in the consumer food industry around sorghum, driven by increased demand for gluten-free, GMO-free plant-based foods.

**Figure 1.**
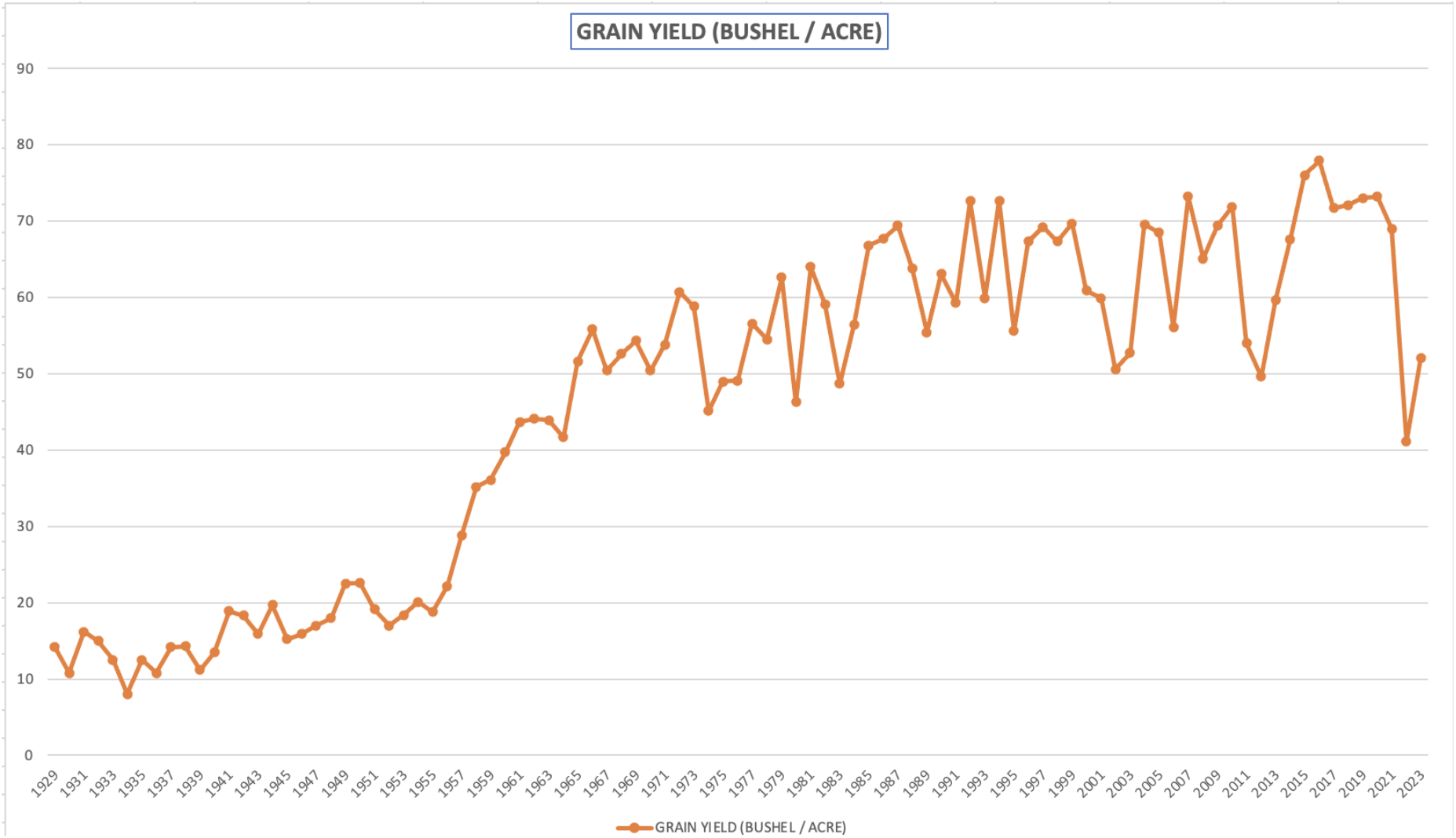
Sorghum yield (bushels per acre) in the US over the last 95 years. Data for this graph was retrieved from USDA National Agricultural Statistics Service (USDA NASS 2025).

Following the paradigm shift to hybrid sorghum production and the conversion of tropical accessions to temperate adaptation, the advent of molecular breeding techniques in the 1990s provided the opportunity for another paradigm shift in sorghum crop improvements. The integration of molecular markers, quantitative trait loci (QTL) mapping, and genomics enabled breeders to identify and manipulate specific genes associated with desirable traits (Mace et al. 2009), with the intent of developing superior sorghum cultivars with improved traits (Xie et al. 2019). In recent decades, genomic selection has been integrated in sorghum breeding to leverage genome-wide markers which predict genetic value and aid in identifying favorable genotypes, thereby accelerating the rate of genetic gain and shortening the breeding cycle (Meuwissen et al. 2001). The bulk of these early inroads in genomic selection utilized genotype-by-sequencing (GBS) to provide the genome-wide markers required. While GBS offers high-density marker discovery and is valuable for *de novo* applications, it often presents challenges related to cost, labor intensity, bioinformatics complexity, and inconsistent data coverage, particularly in resource-limited settings. Furthermore, considerations such as intellectual property rights and commercial access can also limit its broader applicability, particularly in diverse research and applied breeding environments. These limitations are a bottleneck in the use of GBS as a primary marker system for large-scale genomic selection and germplasm characterization in applied breeding programs.

Recognizing the need for a more accessible and cost-effective genotyping array for the US sorghum community, the Crop Germplasm Committee (CGC, https://www.ars-grin.gov/CGC) was tasked in 2022 with establishing a marker resource that meets the needs of geneticists and breeders. The CGC consists of scientists, breeders, and stakeholders focused on the collection, preservation, evaluation, and distribution of germplasm. Subsequently, the sorghum community portal SorghumBase (https://www.sorghumbase.org/; Gladman et al. 2022) coordinated with CGC to establish a CGC genotyping working group with members from academia, government, and industry. The working group was delegated the task to supplement the sorghum molecular breeding toolbox with a new genotyping platform that provides a cost-effective genome-wide marker resource.

Here, we present a community effort to develop an integrated, mid-density, multiplexed, Sorghum Community Genotyping Array (SCGA) designed for various research and breeding applications on the PlexSeq Next Generation Sequencing genotyping platform. This genotyping array represents a cost-effective, high-throughput genetic resource with SNP markers spanning all 10 sorghum chromosomes while also containing genetic markers linked to phenotypic traits of importance to crop improvement and fundamental research programs in sorghum. As a proof-of-concept case study, the sorghum association panel (SAP) (Casa et al. 2008) and a panel of germplasm from public and private sector sorghum breeders were genotyped with the marker array to determine the ability of this platform to capture the genetic diversity residing in these genetic resources. We describe the development of the SCGA and its key features of interest to the sorghum community. Also, within this study, the SCGA was compared to GBS as a genome-wide genotyping assay for use in phylogenetic and genomic selection studies within a diverse collection of sorghum germplasm utilized in basic and applied research programs in the US.

## Materials and Methods

### Development of Sorghum AgriPlex Genomics Mid-density SNP Platform

The CGC genotyping working group conducted surveys of public and private sector researchers to identify their genotyping service needs in support of molecular breeding, diversity studies, and genetic profiling. Using *Sorghum bicolor* v3.1.1 as reference genome of BTx623 (https://phytozome-next.jgi.doe.gov/info/Sbicolor_v3_1_1), a list of 3,496 candidate SNP markers were identified based on existing marker resources for sorghum such as DArTag genotyping service offered by Diversity Arrays Technology Ltd for CGIAR (Consultative Group on International Agricultural Research), community KASP (Kompetitive Allele Specific PCR) panel, and QC (quality control) markers, plus a selection of trait-associated markers identified by breeders (Jaccoud et al. 2001; Kilian et al. 2012; Semagn et al. 2014). Using the list of candidate SNP markers and 150 bp flanking sequences upstream and downstream of the markers, AgriPlex Genomics evaluated each SNP *in silico* using PlexForm™, a proprietary software for primer and multiplex design by AgriPlex Genomics (AgriPlex Genomics, 2025). SNPs shown to be present in multiple locations across the sorghum genome, or in regions of highly repetitive sequences were excluded from the SNP array. Upon the identification of potential informative SNPs, the PlexSeq™ workflow was implemented to develop the platform for the SCGA as follows (Figure 2):

1. The proprietary multiplexing algorithm, PlexForm™ software, is used to design all possible primers around all targeted SNPs. Artificial intelligence algorithms identify the optimal sets of primers that can be mixed in one PCR amplification reaction.
2. Crude DNA isolation from ∼ 50 mg (two leaf punchers) of fresh leaf tissue. The method requires only small quantities of crude DNA that can be isolated from various tissues (although leaf tissue was utilized for the development of the sorghum array).
3. Primary PCR: highly multiplexed, low volume (3ul) PCR amplifications followed by secondary, barcoding PCR amplifications.
4. Once the amplicons are completed, the amplicon mixture is equivalent to barcoded libraries produced from other NGS methods. The process produces amplicon libraries that are equivalent in concentration and do not require additional equalization steps. A mixture of all the libraries is subjected to one bead cleanup and is loaded onto the sequencer.
5. Pooling: barcoded amplicons are combined into one tube, purified, and quantified and subsequently sequenced on an NGS sequencer.
6. Once the sequencing is complete, a proprietary allele frequency-based genotype calling analysis software, PlexCall™, provides an automated sequencer to the data workflow. This Java-based software is tuned for each assay and is fully automated based on only the sequencing output files and a sample sheet indicating sample location on the plate.

**Figure 2.**
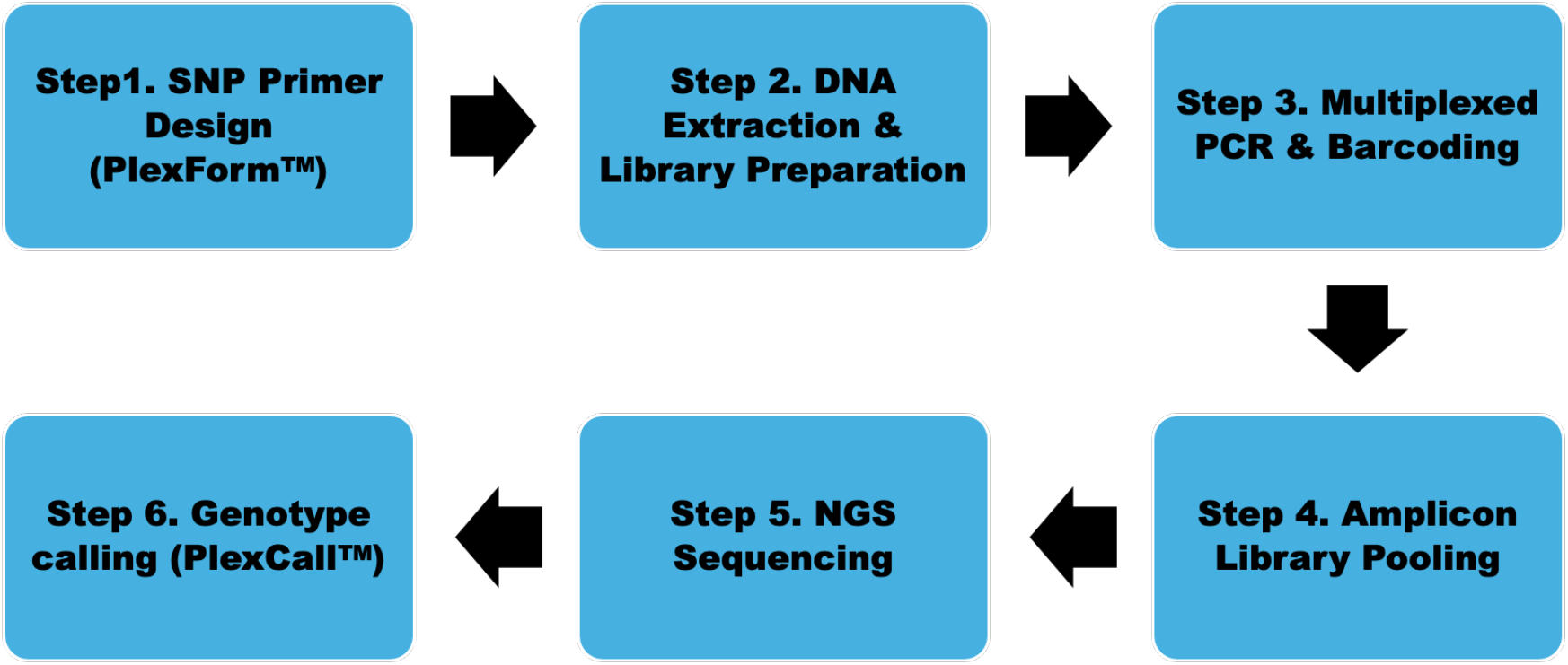
A schematic representation of the PlexSeq™ Next-Generation Sequencing (NGS) genotyping workflow used to develop and implement the Sorghum Community Genotyping Array.

### Germplasm Panel for SNP Array Development

The USDA National Plant Germplasm System (NPGS) conserves a large, diverse set of sorghum germplasm at the Plant Genetic Resources Conservation Unit in Griffin, GA. The collection comprises over 48,500 accessions and includes landraces, cultivars, mapping populations, and sorghum crop wild relatives. There are several defined groups within the collection, including the SAP (Casa et al. 2008), Bioenergy Association Panel (BAP)(Brenton et. al. 2016), and Sorghum Core Collection (Dahlberg et al. 2004). The SAP is highly utilized by the sorghum research community with an average of 1,967 SAP accessions distributed annually by the USDA-ARS Germplasm Resources Information Network (GRIN). Due to its importance to the sorghum research community and access to whole-genome sequencing data (Boatwright et al. 2022), this panel was selected as the NPGS subset used to validate the genotyping platform developed in this project.

The experimental “beta testing” was conducted by genotyping germplasm provided by the USDA NPGS, plus additional samples contributed by more than 15 sorghum-centric programs, including universities, seed companies, and multiple USDA-ARS locations. A total of 2,726 genotypes, including landraces, elite breeding lines, and recombinant inbred lines (RILs), were used to assess the efficacy of the genotyping array to capture a broad range of the genetic diversity in sorghum germplasm. During experimental validation, additional SNPs were removed due to poor performance (Figure 3). The resulting SCGA is made of 2,421 markers, which include 2,365 genome-wide array markers, 26 trait-associated markers (see Supplementary Table), and 30 QC markers.

**Figure 3.**
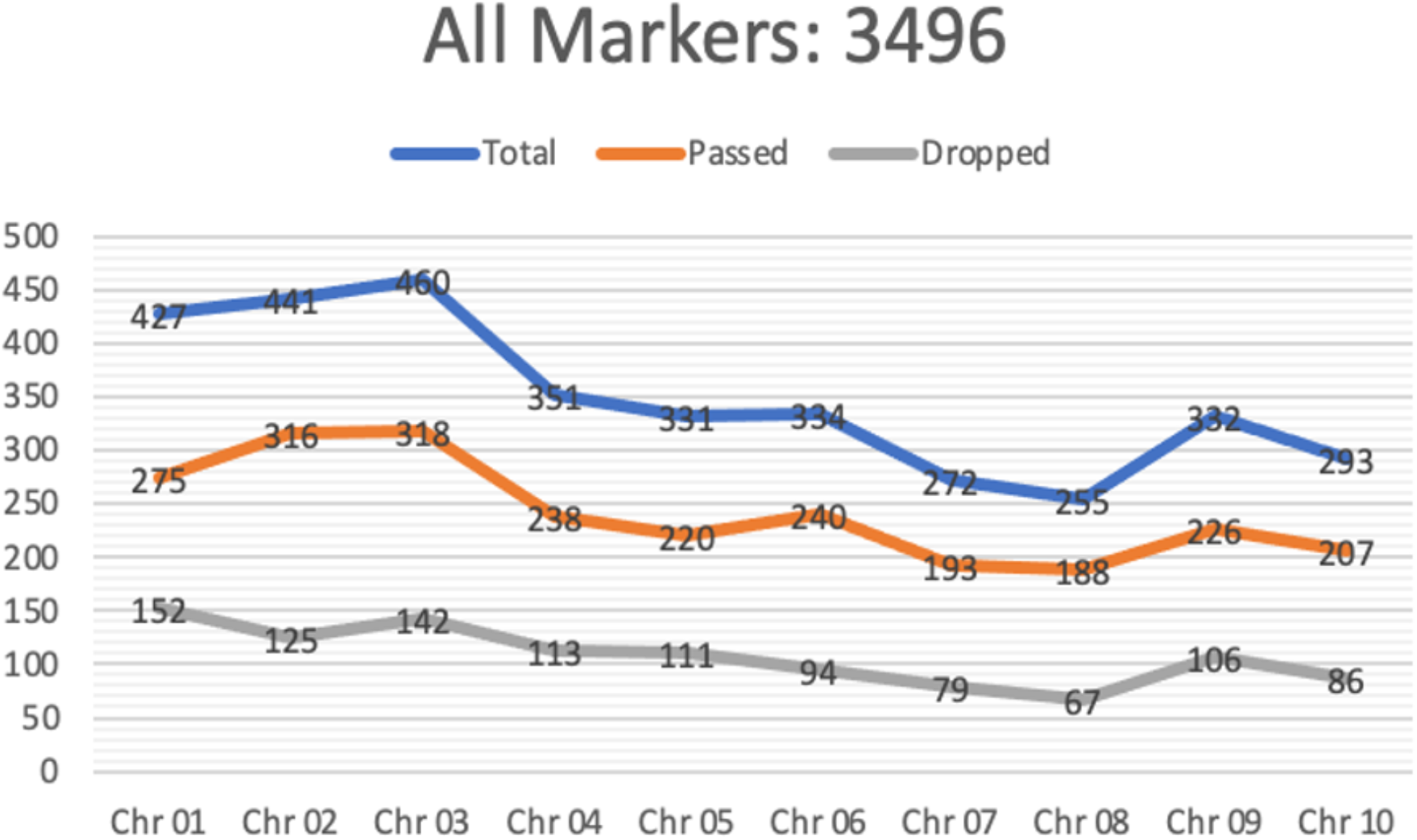
Relative distribution of the total (candidate list), passed (final list), and dropped (eliminated) markers across the ten chromosomes of sorghum.

### Phylogenetic Analyses

A phylogenetic tree was constructed for the SAP population (397 accessions) genotyped using the AgriPlex mid-density sorghum array. The phylogenetic tree was constructed and visualized using Bioconductor R library, ape (Analyses of Phylogenetics and Evolution), version 5.8.1 as an unrooted tree based on the Neighbor-Joining algorithm and the distance matrix.

To compare phylogenetic analysis utilizing Agriplex mid-density array with a high-density GBS array, a germplasm set of 7 seed and 8 pollinator elite inbreds from the Texas A&M AgriLife and Kansas State University sorghum breeding programs were utilized. Using a GBS array with 34,956 SNPs as well as the mid-density sorghum array, principal coordinate analysis with a k-means clustering was conducted (Fonseca et al. 2021; Winans et al. 2023).

### Genomic Selection

To compare genomic selection utilizing Agriplex mid-density array with a high-density GBS array, a set of 56 hybrids from a factorial crossing block of 7 seed and 8 pollinator elite inbreds from the Texas A&M AgriLife and Kansas State University sorghum breeding programs was utilized. These hybrids were grown in 10 environments from South Texas to the High Plains of Kansas to evaluate the following traits: days to mid-anthesis, plant height (cm), and grain yield (standardized to 14% moisture; tons/ha). To determine the utility of the SCGA for use in genomic prediction, models were fit using a high-density GBS array with 34,956 SNPs to compare with models fit using the mid-density SCGA. For parental lines, an additive genomic relationship matrix (GRM) was calculated using a marker matrix coded as A_2_A_2_ = 0, A_1_A_2_ = 1, and A_1_A_1_ = 2 and then mean centered and scaled. For dominance effects, M was recoded as A_2_A_2_ = −2f_i_^2^, A_1_A_2_ = 2f_i_(1-f_i_), and A_1_A_1_ = −2f_i_(1-f_i_)^2^ where f_i_ is the frequency of the rare allele at locus i (Vitezica et al. 2013). GRM were calculated as gBLUP kernels (VanRaden et al. 2008). Hybrid GRM were calculated *in silico* as the Kronecker product between the female and male GRM. The genomic prediction models were:

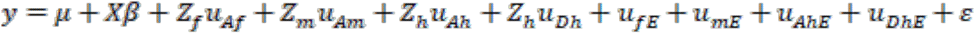

where *y* is the vector of phenotypic values for the trait of interest, *μ* + *Xβ* represents the mean with a fixed effect for environment with incidence matrix *X*, and *ε* represents residual error. Female additive (seed) parent effects are represented by 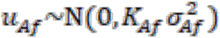 where 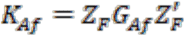 with *G*_*Af*_ as the additive female parental GRM, 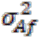 represents the variance component associated with female parent effects. *Z*_*f*_ represents the incidence matrix of female effects. Male additive (pollinator) parent effects are represented by 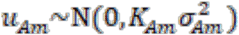 where 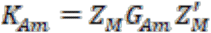 with *G*_*Am*_ as the additive male parental GRM, 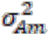 represents the variance component associated with male parent effects. *Z*_*m*_ represents the incidence matrix of male effects. Additive hybrid effects are represented by 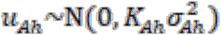 where 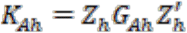 with *G*_*Ah*_ as the additive hybrid GRM, 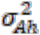 represents the variance component associated with additive hybrid effects, and *Z*_*h*_ represents the incidence matrix of hybrid effects. Dominance hybrid effects are represented by 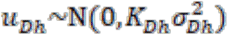 where 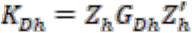 with *G*_*Dh*_ as the dominance hybrid GRM, and 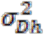 represents the variance component associated with dominance hybrid effects. Interaction effects for males, females, and hybrids (additive and dominance) are represented by 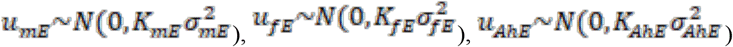, and 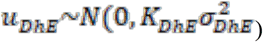, respectively. 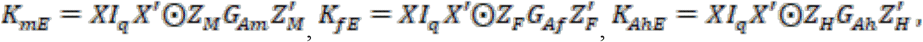 and 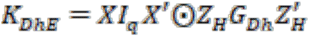 are the kernel structures for the interaction effects with males, females, and hybrids (additive and dominance), respectively. The variance components associated with the interaction effects with males, females, and hybrids (additive and dominance) are represented by 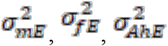, and 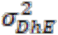, respectively. *I*_*q*_ is an identity matrix for q environments and ⊙ denotes the Hadamard product.

Herein, M1 denotes the model fit using the SCGA and M2 denotes the model fit using the high-density GBS panel. The cross-validation used was a leave-out-line scheme in which each line systematically had all hybrids removed from all environments and were then predicted. Accuracy was determined as the correlation between predicted and actual values for the hybrids of the omitted lines within environments. All models were fitted using the *BGLR* package in R with a Markov Chain Monte Carlo algorithm with 10,000 total iterations, 10% discarded as burn-in, and one in every five samples was retained (thin value of 5).

## Results

The SNPs comprising the SCGA span the 10 sorghum chromosomes with an average of 242 SNPs per chromosome and a range of 188 SNPs (chromosome 8) to 318 (chromosome 3) (Figure 4, Table 1). The genome-wide marker array targeted the gene-rich regions of each chromosome, while markers are, in general, lacking from the gene-poor pericentromeric regions (Figure 4). The number of markers per chromosome is approximately proportional to the relative physical size of the chromosome (Tables 1-2). The distance between adjacent SNPs ranges from 0.23 Mbp to 0.34 Mbp and averages 0.29 Mbp between markers (Table 1; see Supplementary Data File for a complete listing of the markers and their physical positions).

**Table 1.** Number of markers per chromosome and the average physical distance between adjacent SNPs. The 2,421 SNPs are distributed among the 10 sorghum chromosomes with an average of 242 markers per chromosome.

| **Chromosome** | **No. of SNPs** | **Average Distance Between Adjacent SNPs (bp)** |
| --- | --- | --- |
| **1** | 275 | 293,182 |
| **2** | 316 | 243,796 |
| **3** | 318 | 233,499 |
| **4** | 238 | 286,481 |
| **5** | 220 | 324,616 |
| **6** | 240 | 255,249 |
| **7** | 193 | 339,954 |
| **8** | 188 | 333,921 |
| **9** | 226 | 263,156 |
| **10** | 207 | 292,644 |
| **Total** | 2,421 | 286,650 |

**Table 2.**
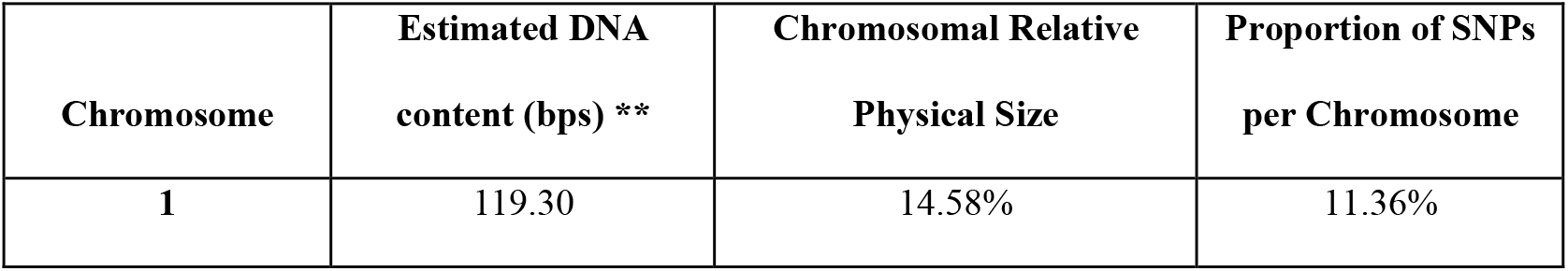
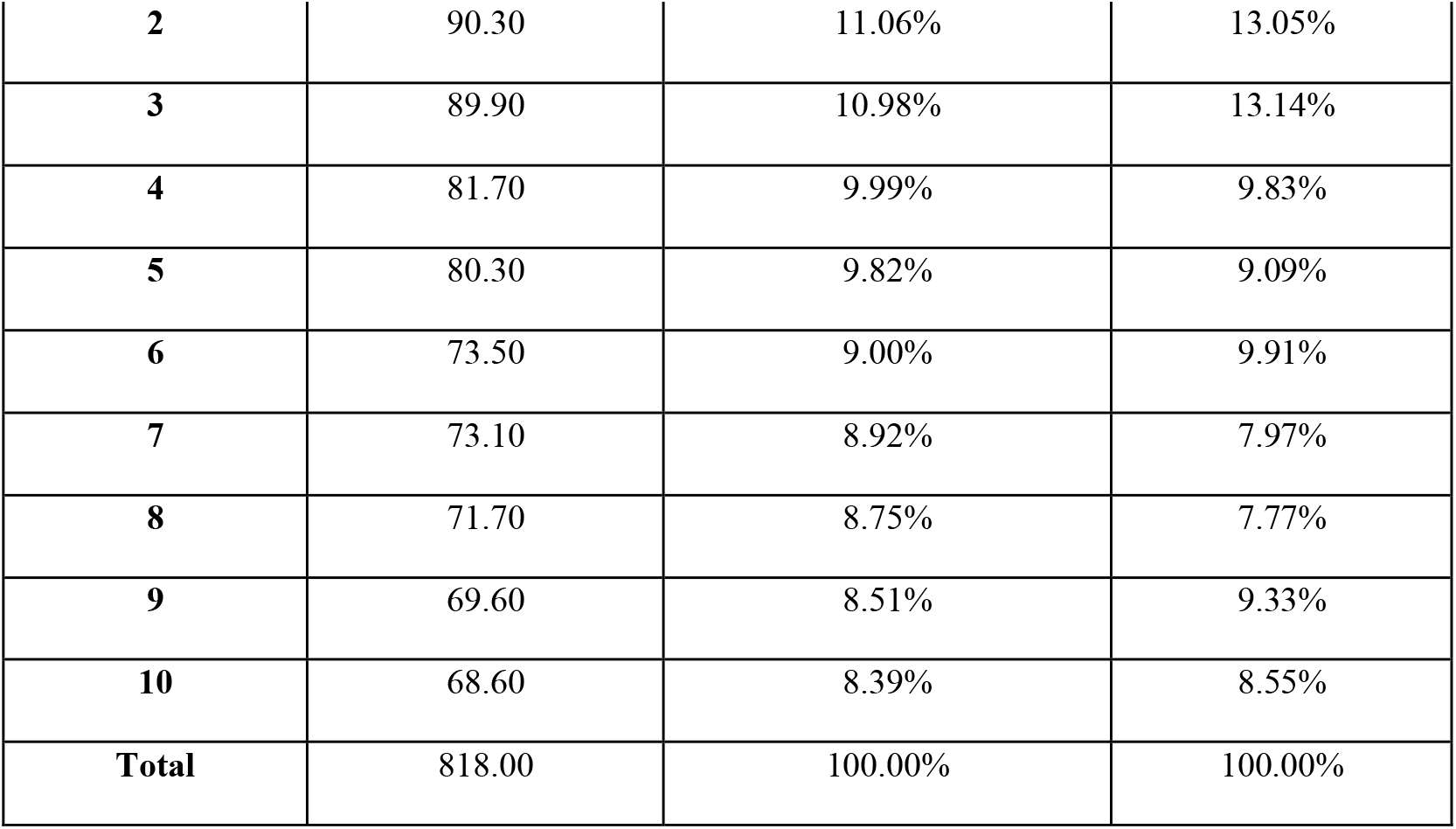
Comparing the proportion of SNPs per chromosome to the relative size of each chromosome. The number of markers per chromosome is relatively proportional to the relative physical size of the chromosomes. **Estimates of chromosome sizes adapted from (Kim et al. 2005).

| **Chromosome** | **Estimated DNA content (bps) \*\*** | **Chromosomal Relative Physical Size** | **Proportion of SNPs per Chromosome** |
| --- | --- | --- | --- |
| **1** | 119.30 | 14.58% | 11.36% |
| **2** | 90.30 | 11.06% | 13.05% |
| **3** | 89.90 | 10.98% | 13.14% |
| **4** | 81.70 | 9.99% | 9.83% |
| **5** | 80.30 | 9.82% | 9.09% |
| **6** | 73.50 | 9.00% | 9.91% |
| **7** | 73.10 | 8.92% | 7.97% |
| **8** | 71.70 | 8.75% | 7.77% |
| **9** | 69.60 | 8.51% | 9.33% |
| **10** | 68.60 | 8.39% | 8.55% |
| **Total** | 818.00 | 100.00% | 100.00% |

**Figure 4.**
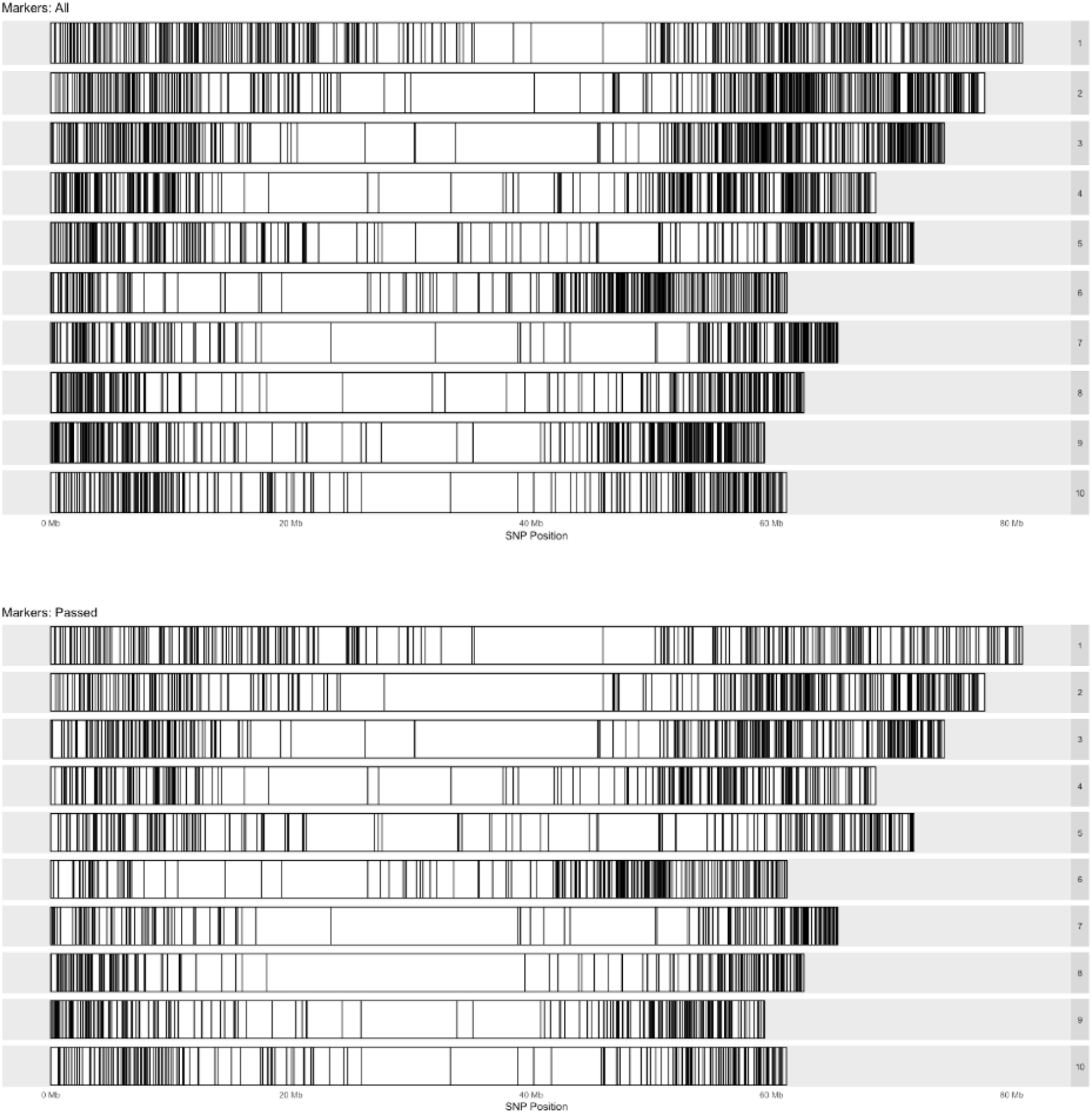
Genome-wide distribution of SNP markers. The top panel illustrates the physical positions of all candidate markers, and the bottom panel shows the final, curated set of markers used in the array. Marker density is visibly lower in the centromeric (gene-sparse) regions of each chromosome, and positions are referenced to the Sorghum bicolor v3.1.1 BTx623 reference genome.

In addition to the genome-wide SNPs within the array are SNPs provisionally linked to traits of agronomic importance to geneticists and sorghum improvement programs (see Supplementary Table). While an exhaustive list of potential traits of interest was not included, this set included resistance to abiotic and biotic stressors and A_1_ CMS fertility restoration loci. It should be noted that the mid-density array was able to differentiate germplasm with resistance or susceptibility traits, such as sorghum aphid resistance and drought tolerance.

### Genetic Characterization of the SAP population using the SNP Array

The sorghum mid-density SNP array was used to genotype existing published populations as a proof of concept of its ability to capture the genetic diversity residing in these genetic resources. Consequently, two different resources were targeted: a large diversity panel (SAP), and a small, targeted breeding panel with germplasm from public and private sector researchers. The SAP population is a collection of converted landraces and temperate-adapted sorghum accessions that represent a broad range of genetic variation within the species, which serves as an excellent germplasm panel for assessing and validating the capabilities of the SCGA.

The mid-density marker array was effective at genotyping the SAP population of 397 accessions (also referred to as samples). A total of 135 (34%) accessions indicated a call rate of 95% or higher, while 387 (97.5%) accessions showed a call rate of 90% or higher. All accessions met a call rate threshold of 60% (Figure 5). Similarly, a total of 1,977 (81.7%) markers were called at 95% or higher and 2,077 (85.8%) at 90% or higher. More than 2,313 (95.5%) markers surpassed a call rate threshold of 60% (Figure 6).

**Figure 5.**
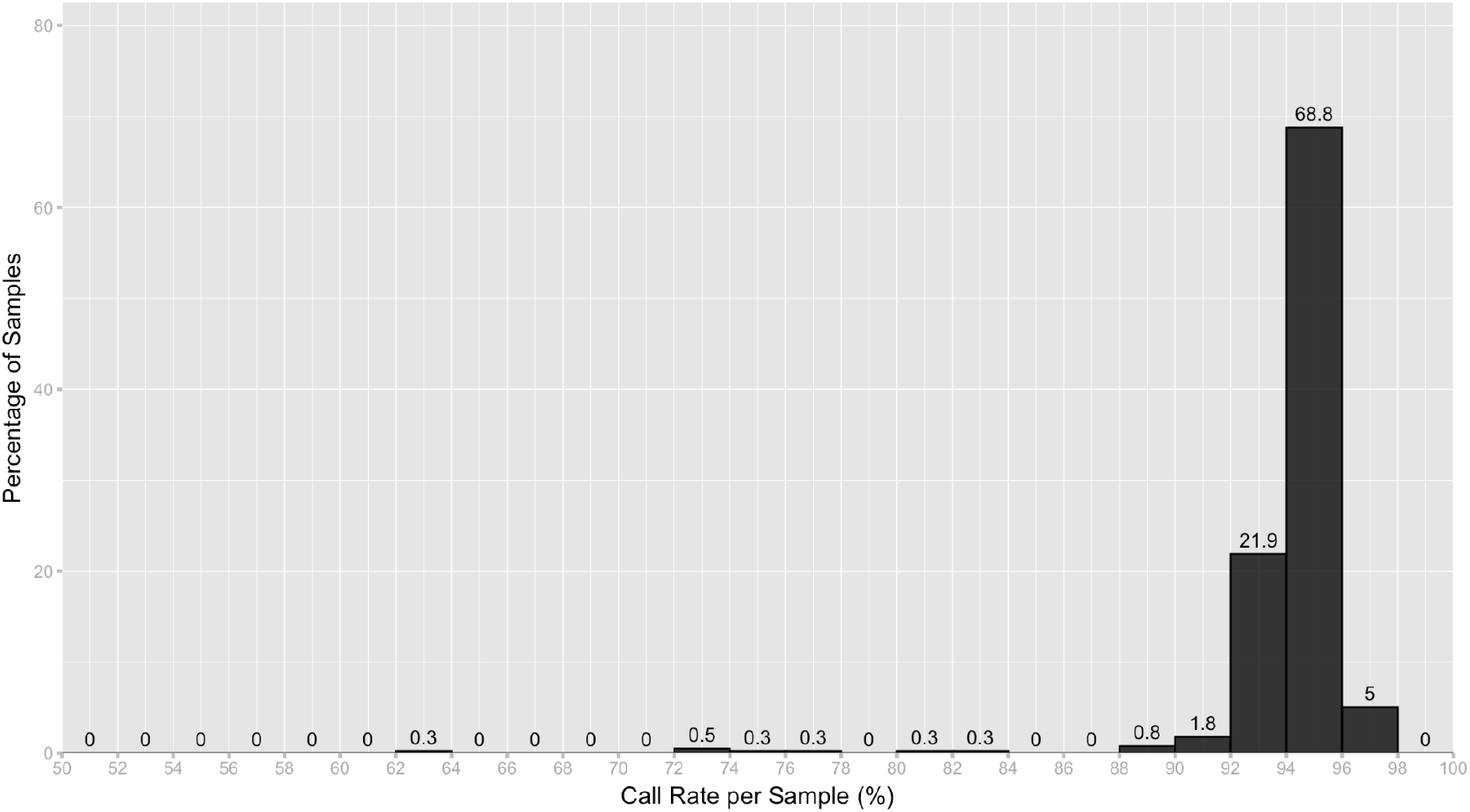
Distribution of call rate per sample based on 2,421 SNPs in SCGA and 397 SAP accessions.

**Figure 6.**
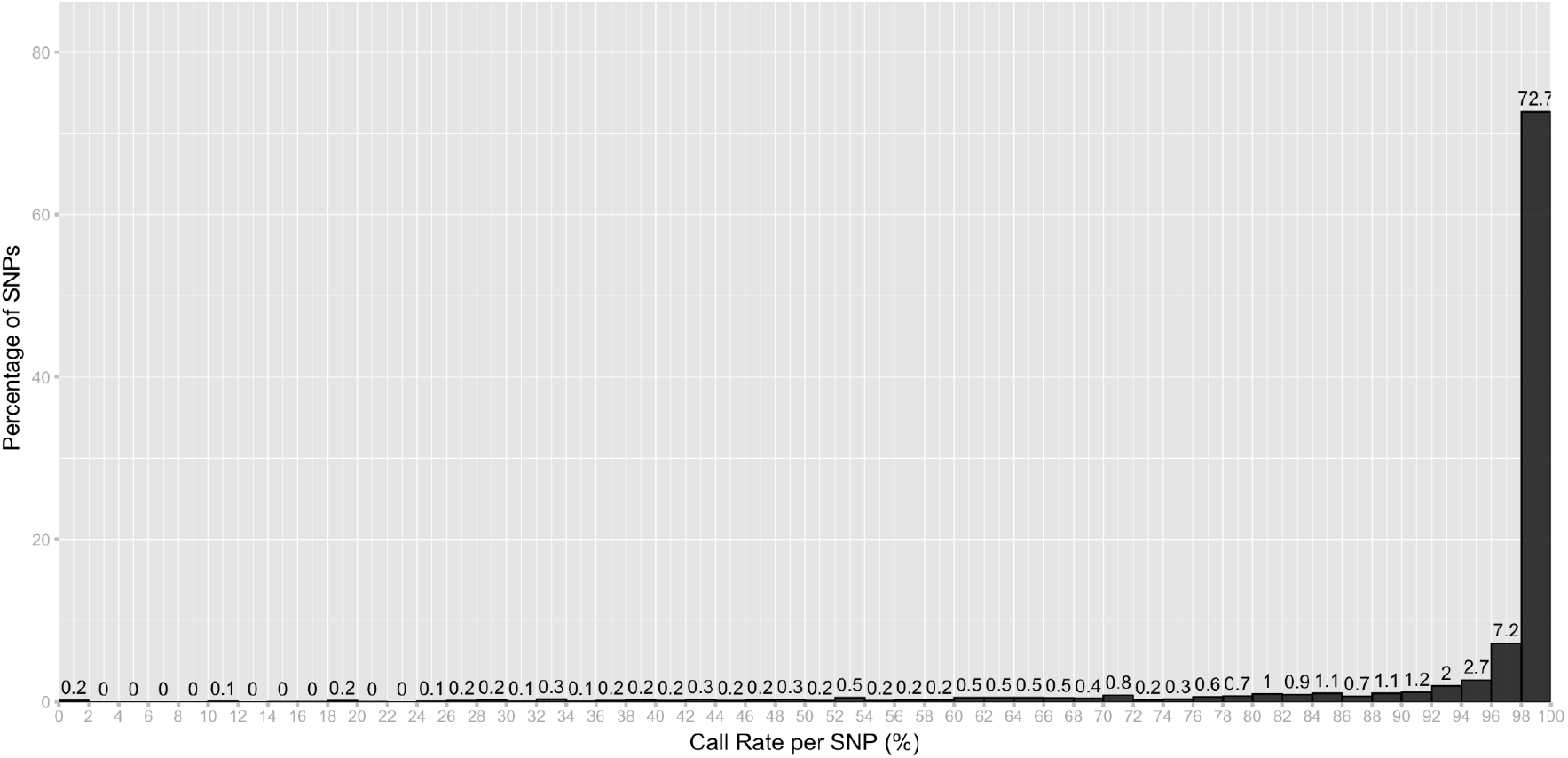
Distribution of call rate per SNP based on 2,421 SNPs in SCGA and 397 SAP accessions.

The SAP panel is predominantly a homozygous population and the SNP array was used to validate heterozygosity across markers and accessions. Of these, 152 (38.3%) accessions were characterized with heterozygosity of 1% or lower, and 394 (99.2%) with heterozygosity of 5% or lower based on the SNP array (Figure 7). Similarly, 2,312 (95.5%) SNPs displayed a heterozygosity of 1% or lower, while 2,350 (97.1%) SNPs indicated a heterozygosity of 5% or lower (Figure 8). Finally, the SAP panel was also assessed for minor allele frequency (MAF). A total of 2,298 (94.9%) SNPs met the MAF threshold of 1%, while 2,266 (93.6%) SNPs had an MAF above 5% (Figure 9).

**Figure 7.**
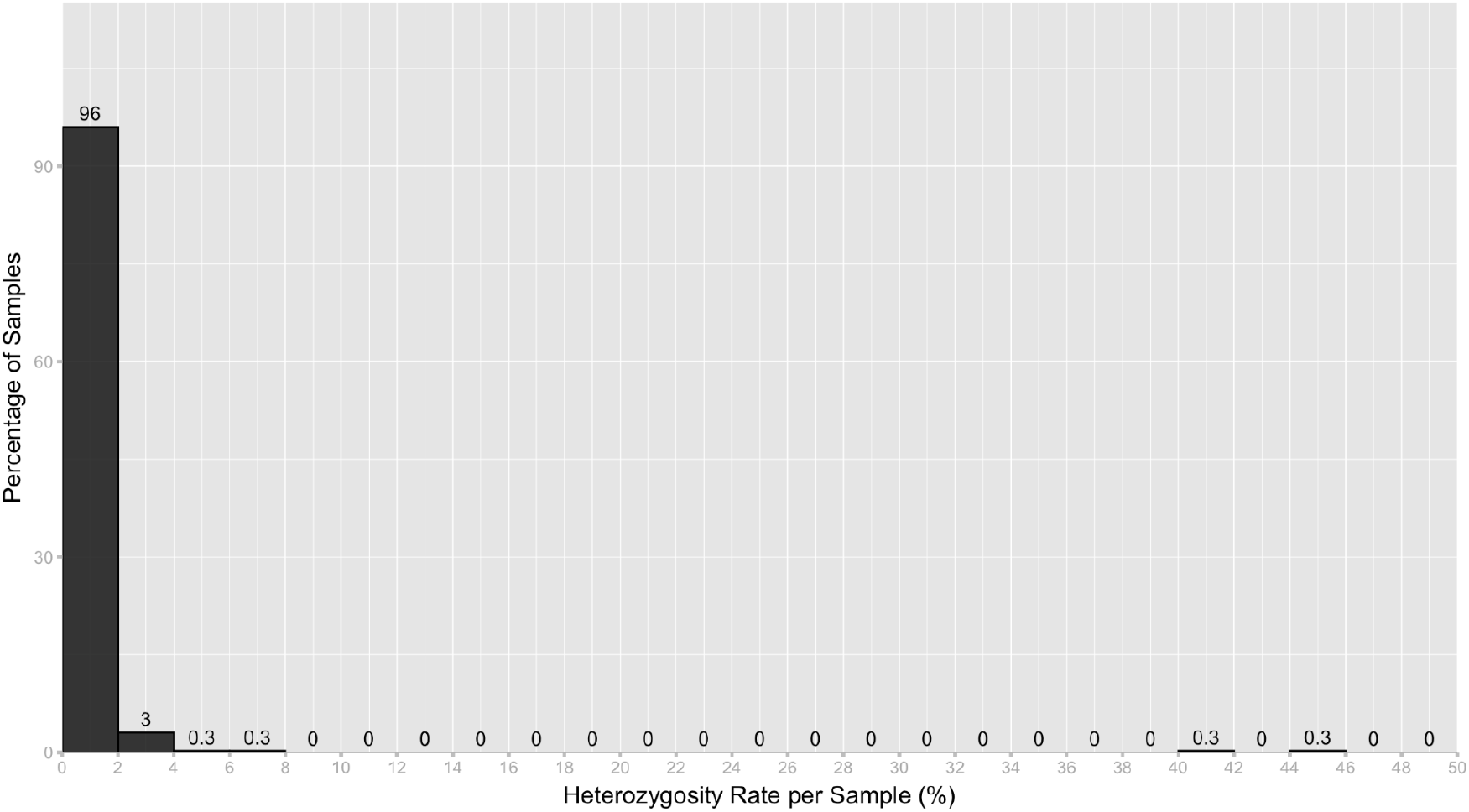
Distribution of heterozygosity rate per sample based on 2,421 SNPs in SCGA and 397 SAP accessions.

**Figure 8.**
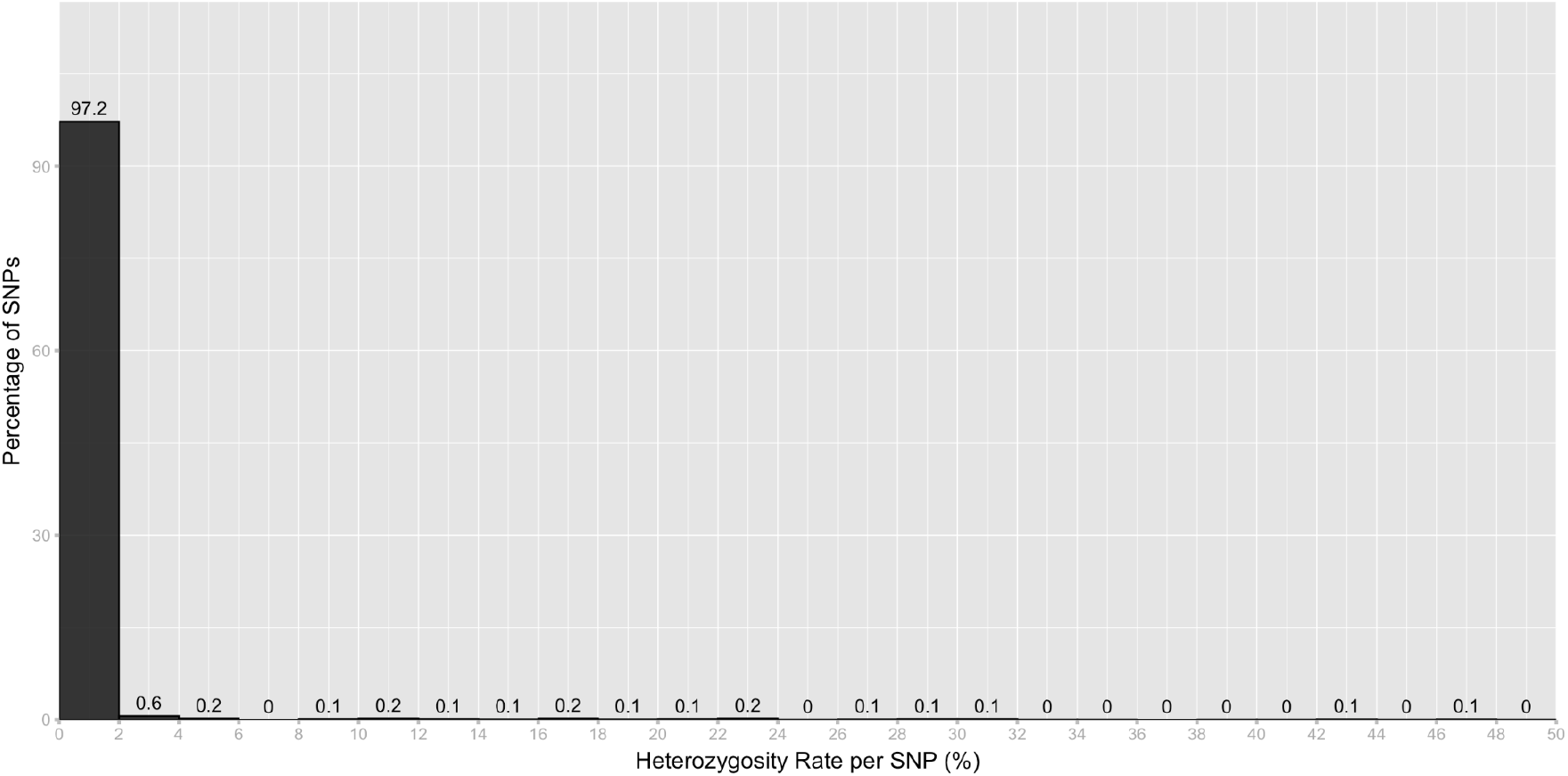
Distribution of heterozygosity rate per SNP based on 2,421 SNPs in SCGA and 397 SAP accessions.

**Figure 9.**
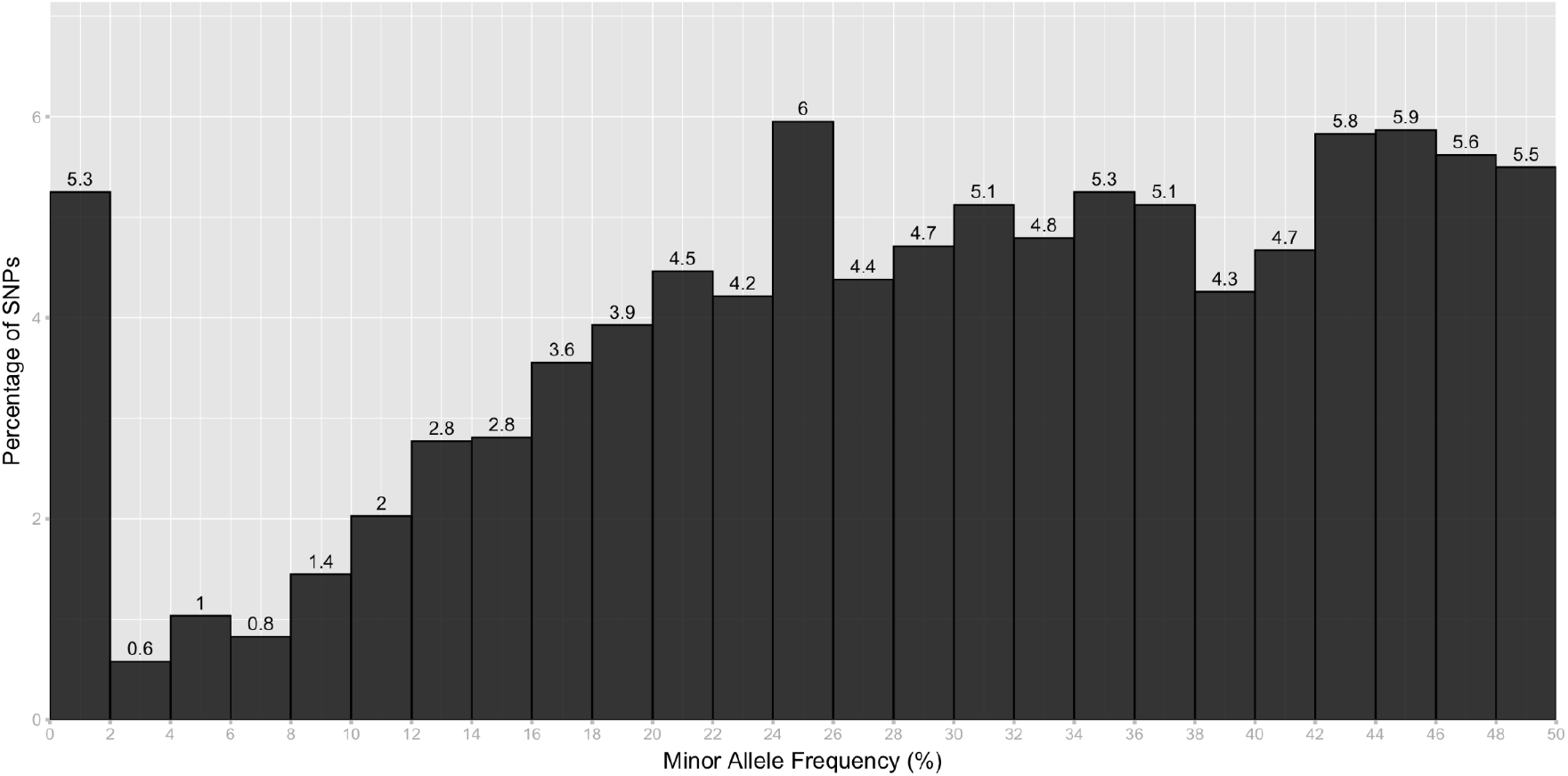
Distribution of minor allele frequency based on 2,421 SNPs in SCGA and 397 SAP accessions.

To investigate the utility of the mid-density array for population structure studies, principal component analysis (PCA) was conducted on 397 SAP accessions genotyped with the marker array. The first ten principal components (PCs) collectively explained 35.5% of the total genetic variance, with PC1 accounting for 12.4%, PC2 for 8.0%, and PC3 for 3.4% (Figure 10). These clustering patterns underscore the array’s ability to resolve major sorghum races within the SAP, even at mid‐density marker coverage.

**Figure 10.**
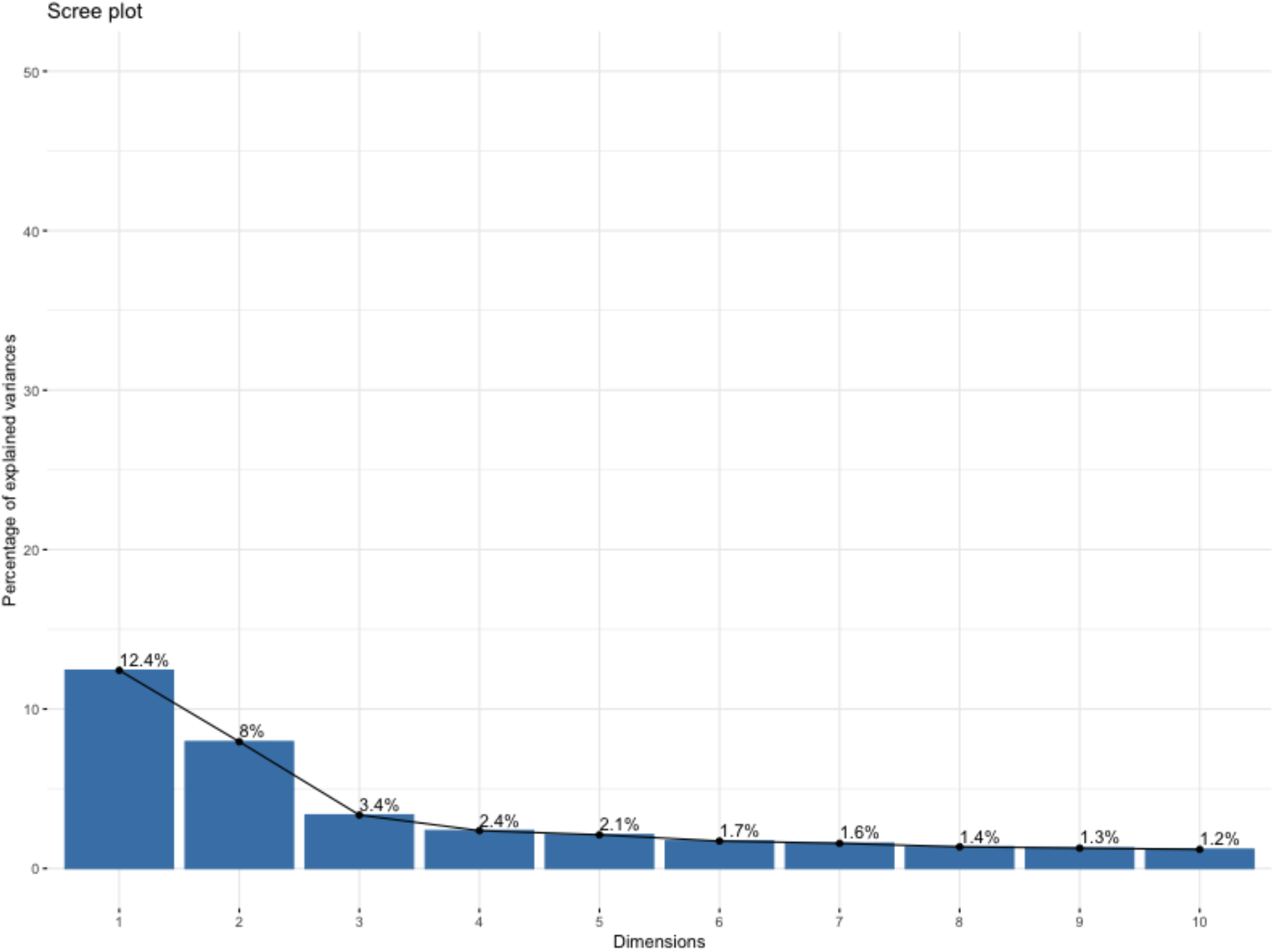
Contribution of the top 10 principal components to the explained variance in the 397 accessions of the SAP panel based on SCGA.

### PCA and Genomic Prediction Model of the Breeding Panel using the SCGA

While the breeding panel examined here consisted of a limited number of elite inbreds, these 15 elite inbreds are representative of a large array of pollinator and seed parental inbreds residing in the Texas A&M and Kansas State University sorghum breeding programs. Furthermore, these parental inbreds were previously included in a genomic prediction study in which a high-density GBS array was employed, which will permit a comparison of model accuracy between the mid-density and high-density marker systems.

The mid-density array performed similarly in the SAP and the breeding panel, for several QC metrics such as call rates and heterozygosity rates computed across 15 samples and 2,421 SNPs (see Supplementary Figures).

The variation explained by the first two principal coordinates for the SCGA was 52.19% while the first two principal coordinates for the high-density marker assay explained 49.88%. While the pollinator parents present in the same k-means cluster did vary between the two marker sets, the clustering and relation of the seed and pollinator parents to each other does appear to be consistent. As a result, the ability to discern the two heterotic pools can be visualized with either marker set (Figures 11 and 12).

**Figure 11.**
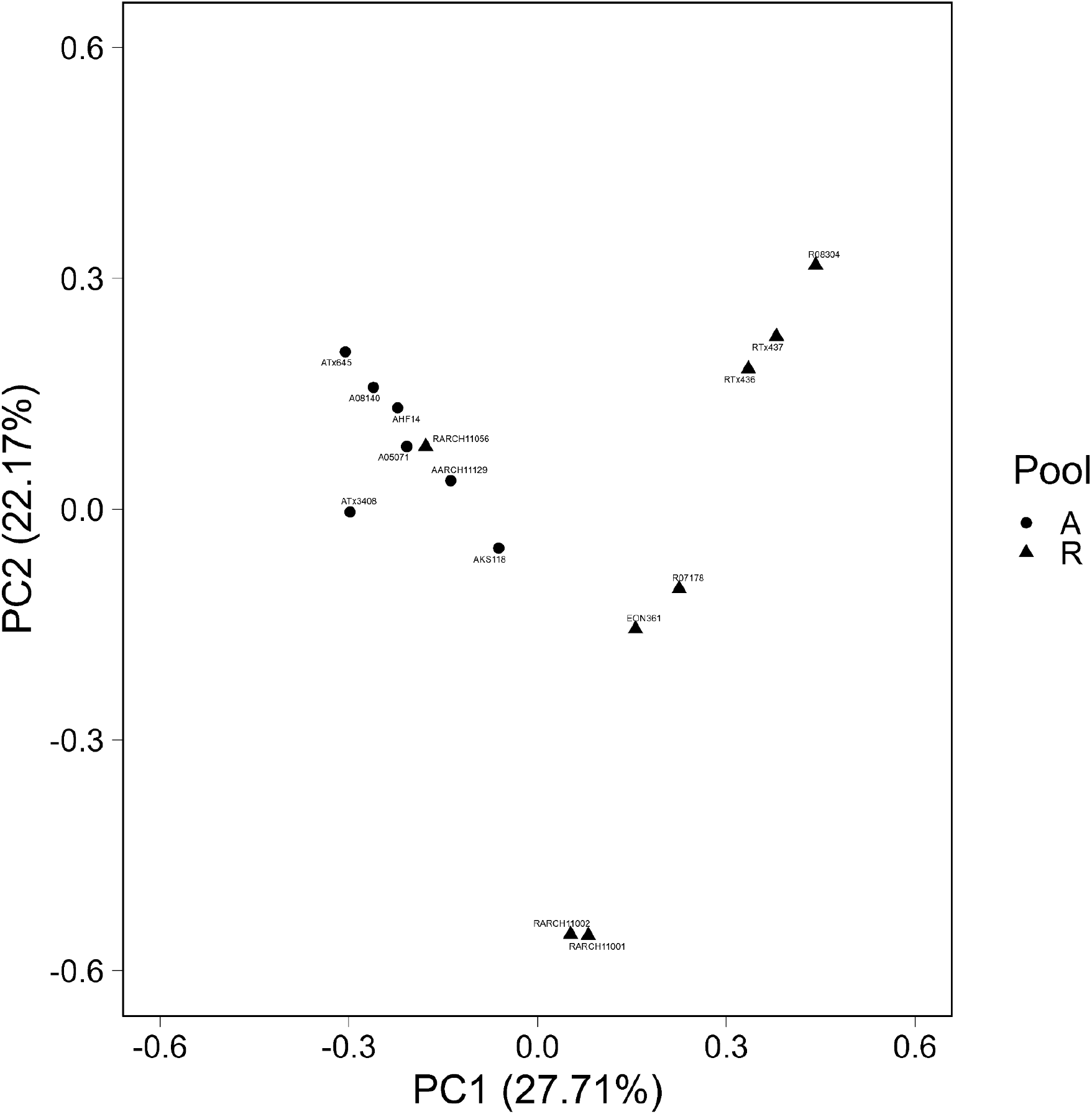
PCA plot based on the GBS platform showing the distribution of the heterotic groups.

**Figure 12.**
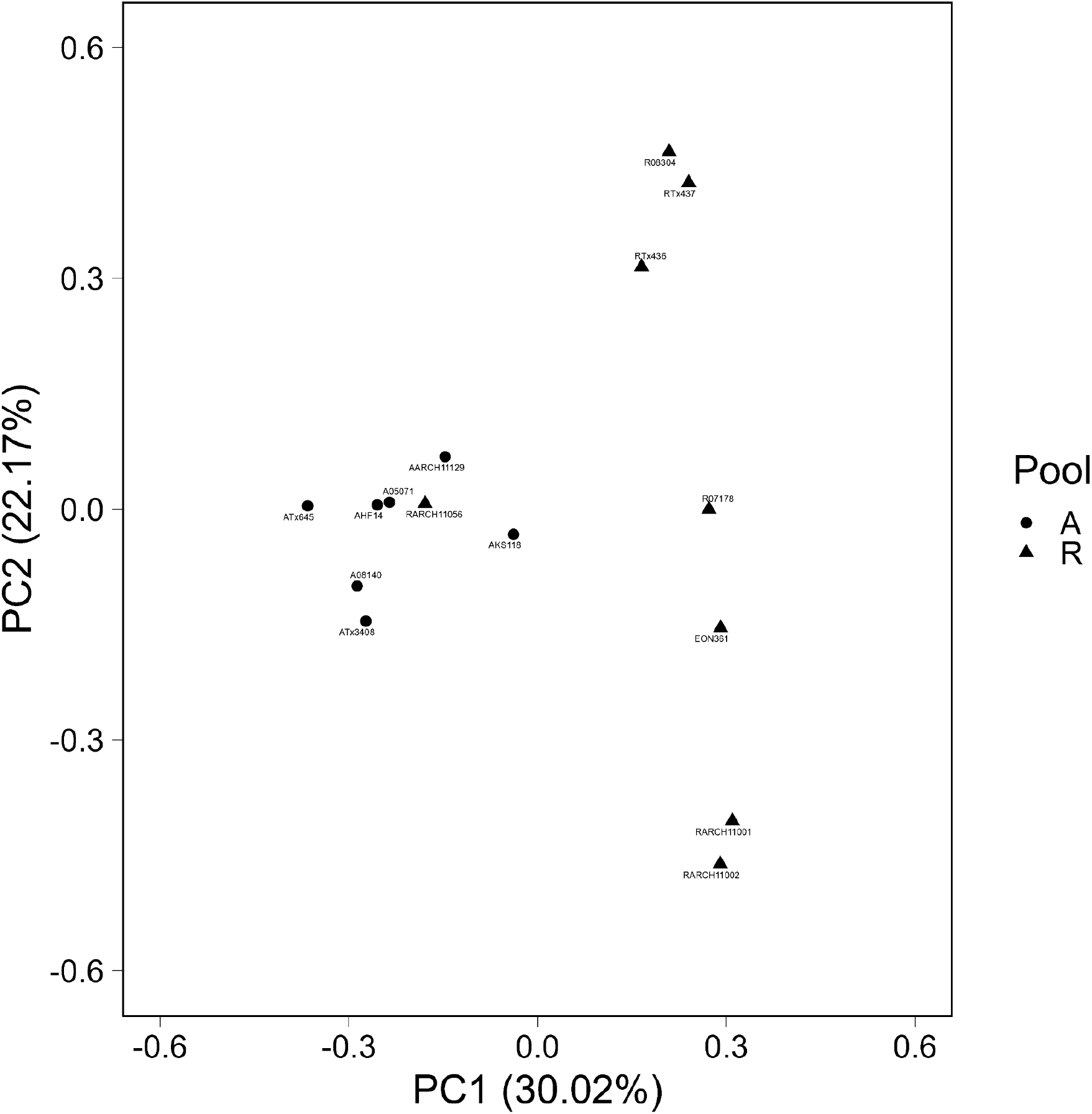
PCA plot based on the mid-density array. Two principal components show a distinct distribution of the heterotic groups, similarly to the one based on GBS.

To allow further comparison of the utility of the mid-density array to a high-density GBS array, genomic prediction models were derived from the mid-density array and compared to published results based on GBS markers. The prediction accuracy for the genomic prediction models using the SCGA were essentially the same as models based on the high-density GBS array. The range of values for both models was the same for grain yield and plant height with correlations of 0.32-0.72 and 0.74-0.88, respectively.

## Discussion

The advancement of genotyping technologies over the past two decades has significantly enhanced the capacity of crop breeding programs to apply molecular tools in the selection and characterization of germplasm. The development and application of a SNP array represent a significant advancement in the genetic tools for characterization of plant germplasm. This study’s primary objective was to develop and evaluate the efficacy of such a marker array in genotyping diverse germplasm, assessing its resolution capabilities, and validating its performance against established marker systems. Our findings demonstrate the mid-density array’s robust performance across a wide range of accessions, providing sufficiently high-resolution genetic data for the identification and introgression of desirable traits into new varieties and various sorghum breeding applications. These results are particularly important in the broader context of germplasm conservation and plant breeding, thus facilitating the development of environmentally resilient, high-yielding, and disease-resistant crops and addressing critical challenges in food security and agricultural sustainability.

### Performance of the Mid-density SNP Array

Among the available platforms, sequencing-based and PCR-based methods have emerged as the most prominent due to their scalability, cost-efficiency, and informativeness. Based on multiplex PCR followed by high-throughput next-generation sequencing (PlexSeq™ NGS workflow), the present array enables the simultaneous genotyping of ∼2,400 user-defined SNP loci per sample at a competitive cost per datapoint. The marker array spans all ten sorghum chromosomes and incorporates trait-linked loci of high relevance to crop improvement for breeders and researchers in the sorghum community.

The performance of the SCGA was rigorously assessed, revealing its high efficiency and accuracy in generating informative genotypic data. Key performance metrics, such as genotyping success rate, call accuracy, and reproducibility, consistently demonstrated the array’s robustness across the diverse germplasm accessions analyzed. As a proof-of-concept, two different populations - a large diversity, research panel in the form of SAP and a small, targeted breeding panel with germplasm from public and private sector researchers - were genotyped with this array to assess its capacity to handle distinct use cases of interest to researchers and breeders.

The high call rates and low missing‐data rates observed in the SAP panel confirm that this array performs reliably across a diverse set of sorghum landraces and temperate-zone lines. Specifically, 97.5% of samples and 85.8% of markers exceeded a 90% call‐rate threshold, demonstrating that the marker system exhibits sample‐ and marker‐level robustness, minimizing downstream imputation and reducing bias in analyses based on allele‐frequency. The extremely low observed heterozygosity (<1% in over 95% of loci) aligns with expected based on the inbreeding levels in the SAP and underscores the array’s ability to discriminate true heterozygotes from background/technical noise. Moreover, with 93.6% of SNPs meeting a minor allele frequency of ≥5%, the SCGA captures ample allele frequency spectrum for diversity analyses, linkage mapping, and genomic prediction. The PCA results (Figure 10) corroborate those of prior whole‐genome sequencing studies of the SAP, which reported that the first ten PCs account for ∼36% of genomic variation, with PC1, PC2, and PC3 explaining 9.36%, 7.86%, and 3.78%, respectively, and similarly partitioned race Kafir, Caudatum/Milo, Durra, and race Guinea (Boatwright et al. 2022). The present mid‐density array captured similar patterns of race differentiation demonstrating its effectiveness in representing core population structure and its potential in studying population stratification, diversity assessment, or correction for structure in genome-wide association and genomic prediction models. Consequently, breeding programs and genebanks can leverage this cost‐efficient array to monitor germplasm structure, inform crossing designs across heterotic groups, and control for subpopulation effects in downstream analyses.

Similarly, the genomic prediction accuracies for key agronomic traits were virtually identical between two platforms; the mid‐density SNP array and the high‐density GBS assay, as the grain yield predictions correlated with observed values at 0.32–0.72, and plant height at 0.74–0.88 for both the Agriplex and GBS datasets. This underscores the value of a carefully designed, targeted genotyping array for breeding applications. Together, the above metrics validate the array as a cost‐effective, mid‐density tool that balances genome coverage with high data quality, which is critical for routine breeding applications and genebank quality assessment alike.

Targeted primarily for the U.S. sorghum community, the SCGA offers several advantages for all crop improvement programs including grain, forage, and bioenergy. First, it draws on a broad consensus of public and private sector needs, ensuring that the genotyping array includes markers relevant to stay-green, drought tolerance, Striga resistance, and other priority traits (Zebire et al. 2021, Cuevas et al. 2022). Second, by standardizing on a mid-density, sequencing-based genotyping workflow, breeders can more readily share data across programs, accelerate recurrent selection cycles, and integrate genomic predictions into decision-making to support accelerated breeding. Third, the community-driven genotyping array can be developed and updated in an iterative and agile manner. As new functional variants are discovered, additional loci can be incorporated without changing chemistry, thus preserving backward compatibility and investment in assay development.

### SCGA and the Utility for the U.S. National Plant Germplasm System

We developed this array in coordination with our collaborators at the U.S. NPGS with the goal of further supporting its operations and users. Administered by USDA’s Agricultural Research Service, NPGS aims to conserve the genetic diversity of agriculturally important plants while facilitating the use of germplasm for research, breeding, and education through its GRIN-Global database. The GRIN-Global system curates and distributes thousands of sorghum accessions annually to researchers and breeders, and the ability to efficiently genotype these accessions with a standardized, mid-density SNP array would greatly enhance the value, transparency, and reproducibility of the germplasm it provides. Existing genotypic data for sorghum accessions available through GRIN-Global is largely generated from genome-wide SNP discovery platforms like GBS or DArTseq. While these platforms are valuable for initial characterization and diversity studies, their high rates of missing data and inconsistent coverage across accessions pose challenges for germplasm users seeking to compare or replicate results across downstream studies. In addition, breeders and researchers encounter routine quality control issues resulting from labeling errors, seed mixture events, and unintended cross-pollination. A community-validated and reproducible mid-density SNP array would enable GRIN-Global and its users to genotype existing and regenerated accessions with consistent, high-quality data, facilitating identity verification, seed/germplasm purity testing, and passport data enrichment. For predominantly inbred line crops like sorghum, which are expected to remain genetically stable over cycles, a cost-effective and easily accessible genotyping with this SNP array would further ensure the genetic fidelity and identity of distributed material.

Having an inexpensive, high throughput platform to genotype the entire sorghum collection will provide many benefits to curation of the collection including core collection validation, quality control during regenerations, and identification of redundant accessions. The current USDA sorghum core collection was established based on passport and descriptor data (Dahlberg et al. 2004). Ideally, a core collection should be approximately ten percent of the entire collection and represent the overall diversity of the entire collection (Brown 1989). Since passport data is limited for much of the overall sorghum collection, the ability to use this data to capture the diversity of the entire collection is also limited. Using genetic markers to establish the core or validate the diversity contained in the current core is needed and now possible. The more representative the core collection is of the entire collection, the more beneficial it will be to plant breeders. Genotyping of the collection will also provide curators with a means for quality control of regeneration. This includes verification of accessions at harvest and monitoring of genetic drift after several regeneration cycles. Large germplasm collections like sorghum often have an increased chance for redundant accessions. An effort was made in the past by curators to identify redundancies in the USDA sorghum collection using passport data focused particularly on country of origin and secondary identifiers such as ICRISAT numbers. Again since passport data is limited, it is probable that using passport data alone will miss a significant portion of redundant accessions. Genotyping has been shown to successfully identify redundancies in germplasm collections (Gross et al. 2012; Anglin et al. 2025) and should prove successful for the USDA sorghum collection. Identification of redundancies will allow curators to prioritize regenerations and become more efficient in their regeneration efforts. It will also save users of the germplasm collection from phenotyping accessions that are the same or very similar.

## Conclusions

This study successfully demonstrates the development and comprehensive evaluation of a cost-effective, mid-density SNP array, establishing its utility as a vital genotyping platform for sorghum researchers and breeders. Our findings confirm that this targeted, multiplexed genotyping array, built on the Agriplex Genomics PlexSeq™ platform, effectively captures sufficient polymorphism and provides robust genome coverage across all ten chromosomes of *Sorghum bicolor*. The array’s high call rates, low missing data, and accurate representation of genetic diversity, as evidenced by its ability to resolve major sorghum races within the SAP population consistent with prior whole-genome sequencing studies (Boatwright et al. 2022), underscore its reliability. Crucially, the genomic prediction accuracies achieved with this mid-density array for key agronomic traits were found to be comparable to those obtained using a high-density GBS assay, highlighting its efficiency and practical value for breeding applications (Fonseca et al. 2021). Overall, our results emphasize the significant contribution of the Agriplex Genomics SNP array as a tailored genotyping solution for the U.S. sorghum community. By integrating user-driven marker selection with high multiplexing capacity and sequencing-based accuracy, the platform delivers reproducible genotype calls with minimal missing data and rapid turnaround times. These attributes make it ideally suited for a range of routine applications, including marker-assisted selection (MAS), background recovery in backcross programs, genetic purity testing, and essential germplasm quality control within both breeding pipelines and genebank service providers like the USDA NPGS and its GRIN-Global database. The flexible design of this array further ensures its long-term relevance, allowing for seamless integration of newly discovered trait-linked markers as breeding priorities evolve. This adaptability will sustain its utility in accelerating genetic gain across diverse sorghum improvement efforts, ultimately contributing to enhanced crop resilience and global food security.

## References

AgriPlex Genomics. (2025). What is PlexSeq. Retrieved from https://www.agriplexgenomics.com/plexseq-technology

Anglin NL, Wenzl P, Azevedo V, Lusty C, Ellis D, Gao D (2025) Genotyping genebank collections: Strategic approaches and considerations for optimal collection management. Plants 14(2):252. 10.3390/plants14020252

Boatwright JL, Sapkota S, Jin H, Schnable PS, Morris GP (2022) Sorghum Association Panel whole-genome sequencing establishes cornerstone resource for dissecting genomic diversity. Plant J 111:888–904. 10.1111/tpj.15853

Brenton ZW, Cooper EA, Myers MT, Boyles RE, Shakoor N, Zielinski KJ, … Kresovich S (2016) A genomic resource for the development, improvement, and exploitation of sorghum for bioenergy. Genetics 204:21–33. 10.1534/genetics.115.183947

Brown AHD (1989) Core collections: A practical approach to genetic resources management. Genome 31:818–824. 10.1139/g89-144

Casa AM, Pressoir G, Brown PJ, Mitchell SE, Rooney WL, Tuinstra MR, … Kresovich S (2008) Community resources and strategies for association mapping in sorghum. Crop Sci 48:30–40. 10.2135/cropsci2007.02.0080

Cuevas HE, Knoll JE, Harris-Shultz KR, Punnuri SM (2022) Genetic mapping of sugarcane aphid resistance in sorghum line SC112-14. Crop Sci 62:2267–2275. 10.1002/csc2.20818

Dahlberg JA, Burke JJ, Rosenow DT (2004) Development of a Sorghum Core Collection: Refinement and evaluation of a subset from Sudan. Economic Botany 58:556–567. 10.1663/0013-0001(2004)058[0556:DOASCC]2.0.CO;2

Duncan RR, Bramel-Cox PJ, Miller FR (1991). Contributions of Introduced Sorghum Germplasm to Hybrid Development in the USA. In Use of Plant Introductions in Cultivar Development Part 1 (eds H.L. Shands and L.E. Wiesner). 10.2135/cssaspecpub17.c5

Fonseca JMO, Klein PE, Crossa J, Rooney WL, Stojšin D, Bhosale S, et al (2021) Assessing combining abilities, genomic data, and genotype × environment interactions to predict hybrid grain sorghum performance. Plant Genome 14:e20127. 10.1002/tpg2.20127

FAO (Food and Agriculture Organization of the United Nations) (2022) The State of Food and Agriculture 2022: Leveraging agricultural automation for transforming agrifood systems. Rome: FAO. 10.4060/cb9479en

FAO (Food and Agriculture Organization of the United Nations) (2021) The State of Food and Agriculture 2021: Making agrifood systems more resilient to shocks and stresses. Rome: FAO. 10.4060/cb4476en

Gladman N, Olson A, Wei S, Chougule K, Lu Z, Tello-Ruiz M, Meijs I, Van Buren P, Jiao Y, Wang B, Kumar V, Kumari S, Zhang L, … Ware D (2022). SorghumBase: a web-based portal for sorghum genetic information and community advancement. Planta, 255(2), 35. 10.1007/s00425-022-03821-6

Gross BL, Volk GM, Richards CM, Forsline PL, Fazio G, Aldwinckle HS, et al (2012) Identification of “duplicate” accessions within the USDA-ARS National Plant Germplasm System Malus collection. J Am Soc Hortic Sci 137:333–342. 10.21273/JASHS.137.5.333

Jaccoud D, Peng K, Feinstein D, Kilian A (2001) Diversity Arrays: A solid state technology for sequence information independent genotyping. Nucleic Acids Res 29:e25. 10.1093/nar/29.4.e25

Khalifa M, Eltahir EAB (2023) Assessment of global sorghum production, tolerance, and climate risk. Front Sustain Food Syst 7:1184373. 10.3389/fsufs.2023.1184373

Kilian A, Wenzl P, Huttner E, Carling J, Xia L, Blois H, et al (2012) Diversity Arrays Technology: A generic genome profiling technology on open platforms. Methods Mol Biol 888:67–89. 10.1007/978-1-61779-870-2_5

Kim J-S, Islam-Faridi MN, Klein PE, Stelly DM, Price HJ (2005) Comprehensive molecular cytogenetic analysis of sorghum genome architecture: Distribution of euchromatin, heterochromatin, genes, and recombination in comparison to rice. Genetics 171:1963–1976. 10.1534/genetics.105.048215

Mace ES, Rami J-F, Bouchet S, Klein PE, Klein RR, Kilian A, et al (2009) A consensus genetic map of sorghum that integrates multiple component maps and high-throughput Diversity Array Technology (DArT) markers. BMC Plant Biol 9:13. 10.1186/1471-2229-9-13

Mann JA, Kimber CT, Miller FR (1983) The origin and early cultivation of sorghums in Africa. Texas Agricultural Experiment Station Bulletin No. 1454, College Station, TX. Retrieved from https://oaktrust.library.tamu.edu/handle/1969.1/128074

McCollough RL (1972). Nutritive value of eight hybrid sorghum grains and three hybrid corns compared in all-concentrate rations, Part I: Hybrid sorghum and corn characteristics and methods used to nutritionally evaluate them,” Kansas Agricultural Experiment Station Research Reports: Vol. 0: Iss. 1. 10.4148/2378-5977.2796

Meuwissen THE, Hayes BJ, Goddard ME (2001) Prediction of total genetic value using genome-wide dense marker maps. Genetics 157:1819–1829. 10.1093/genetics/157.4.1819

Mwamahonje A, Mdindikasi Z, Mchau D, Ngwira P, Mneney E, Mutegi E, et al (2024) Advances in sorghum improvement for climate resilience in the global arid and semi-arid tropics: A review. Agronomy 14:3025. 10.3390/agronomy14123025

Semagn K, Babu R, Hearne S, Olsen M (2014) Single nucleotide polymorphism genotyping using Kompetitive Allele Specific PCR (KASP): Overview of the technology and its application in crop improvement. Mol Breed 33:1– 14. 10.1007/s11032-013-9917-x

Tanwar R, Panghal A, Chaudhary G, Kumari A, Chhikara N (2023) Nutritional, Phytochemical and Functional Potential of Sorghum: A Review. Food Chemistry Advances, 3, Article 100501. 10.1016/j.focha.2023.100501

USDA NASS (2025) Data & Statistics. Washington, DC: U.S. Department of Agriculture, National Agricultural Statistics Service. Retrieved from: https://www.nass.usda.gov/Data_and_Statistics/index.php

VanRaden PM (2008) Efficient methods to compute genomic predictions. J Dairy Sci 91:4414–4423. 10.3168/jds.2007-0980

Vitezica ZG, Varona L, Legarra A (2013) On the additive and dominant variance and covariance of individuals within the genomic selection scope. Genetics 195:1223–1230. 10.1534/genetics.113.155176

Whole Grains Council. (2025). Sorghum: June Grain of the Month. Retrieved from https://wholegrainscouncil.org/whole-grains-101/grain-month-calendar/sorghum-june-grain-month

Winans ND, Klein RR, Fonseca JMO, Klein PE, Rooney WL (2023) Evaluating introgression sorghum germplasm selected at the population level while exploring genomic resources as a screening method. Plants 12:444. 10.3390/plants12030444

Xie Q, Xu Z (2019) Sustainable Agriculture: From Sweet Sorghum Planting and Ensiling to Ruminant Feeding. Molecular Plant 12(5):603–606. 10.1016/j.molp.2019.04.001

Zebire D, Menkir A, Adetimirin V, Mengesha WA, Meseka SK, Oyekunle M, et al (2021) Identifying suitable tester for evaluating Striga resistant maize inbred lines using DArTseq markers and agronomic traits. PLoS One 16:e0253481. 10.1371/journal.pone.0253481

