## Supplemental Figures for "Development and evaluation of a cost-effective, mid-density SNP array as a sorghum community genotyping resource"

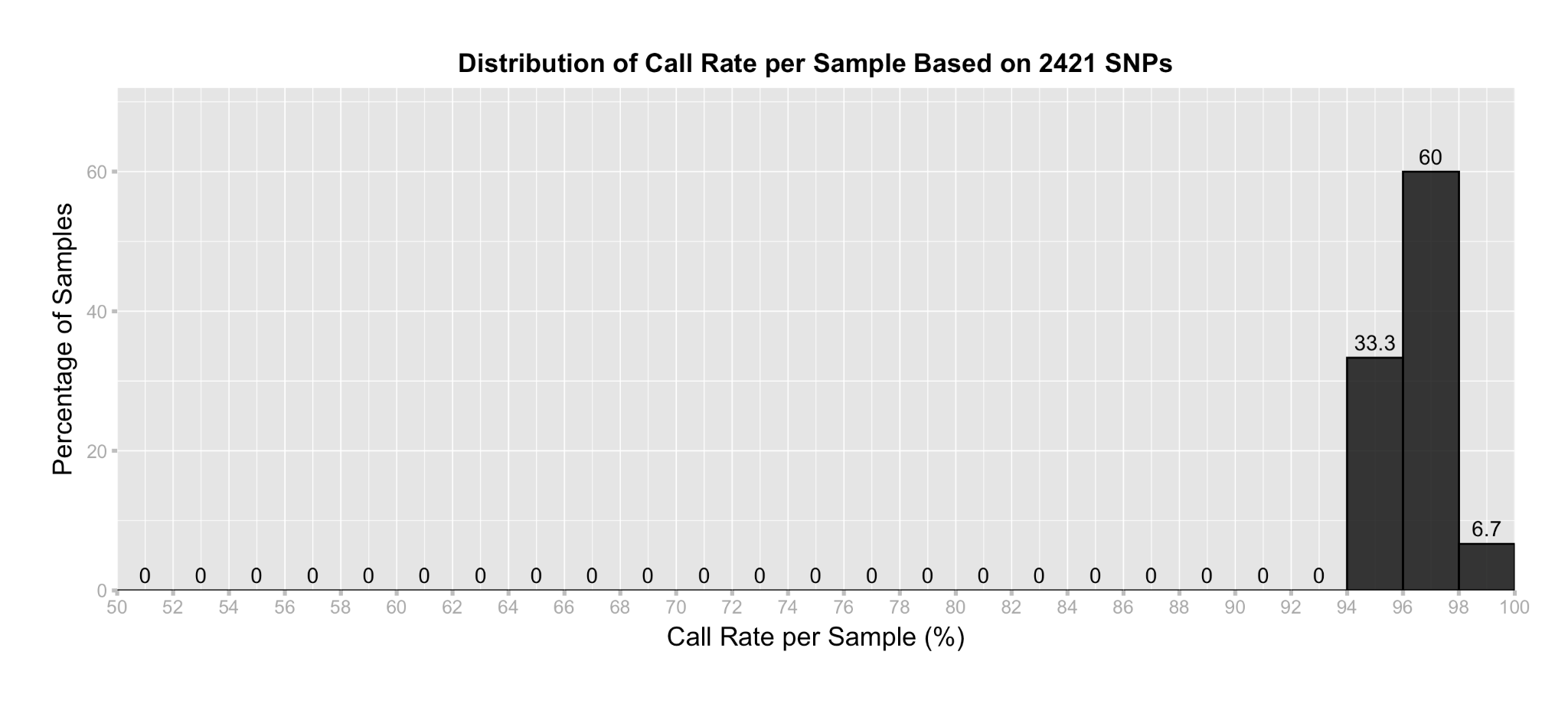

**Supplementary Figure 1.** Distribution of call rate per sample based on 2421 SNPs in SCGA and 15 breeding accessions.

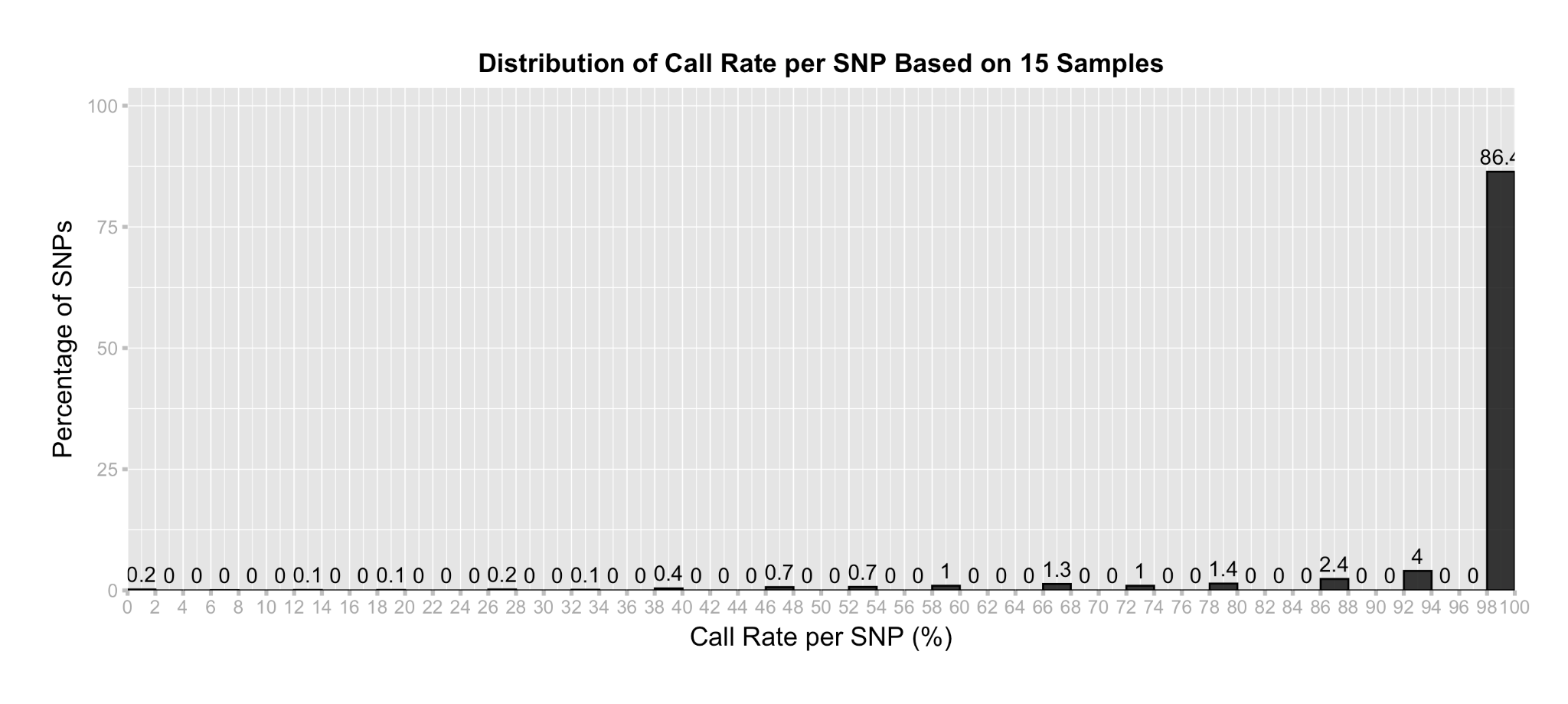

**Supplementary Figure 2.** Distribution of call rate per SNP based on 2421 SNPs in SCGA and 15 breeding accessions.

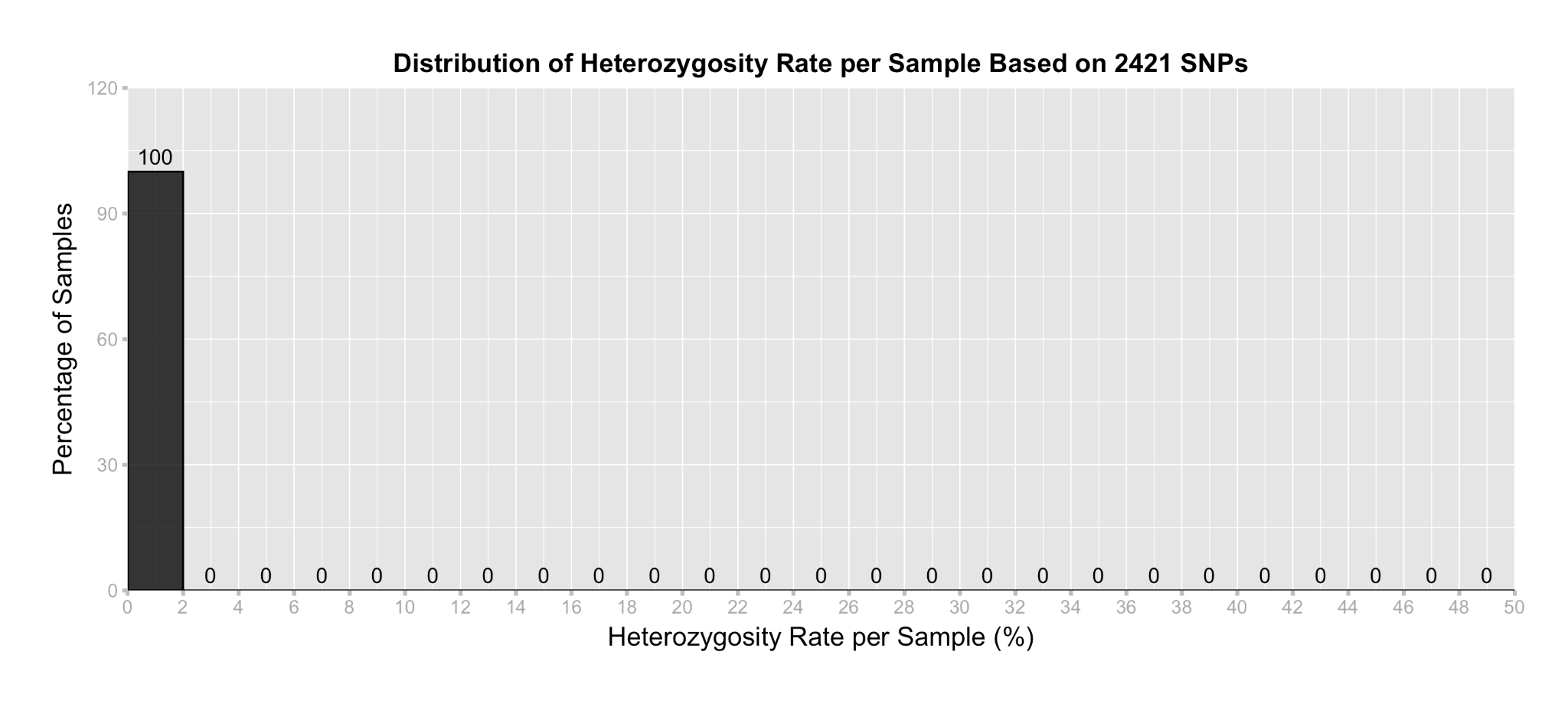

**Supplementary Figure 3.** Distribution of heterozygosity rate per sample based on 2,421 SNPs in SCGA and 15 breeding accessions.

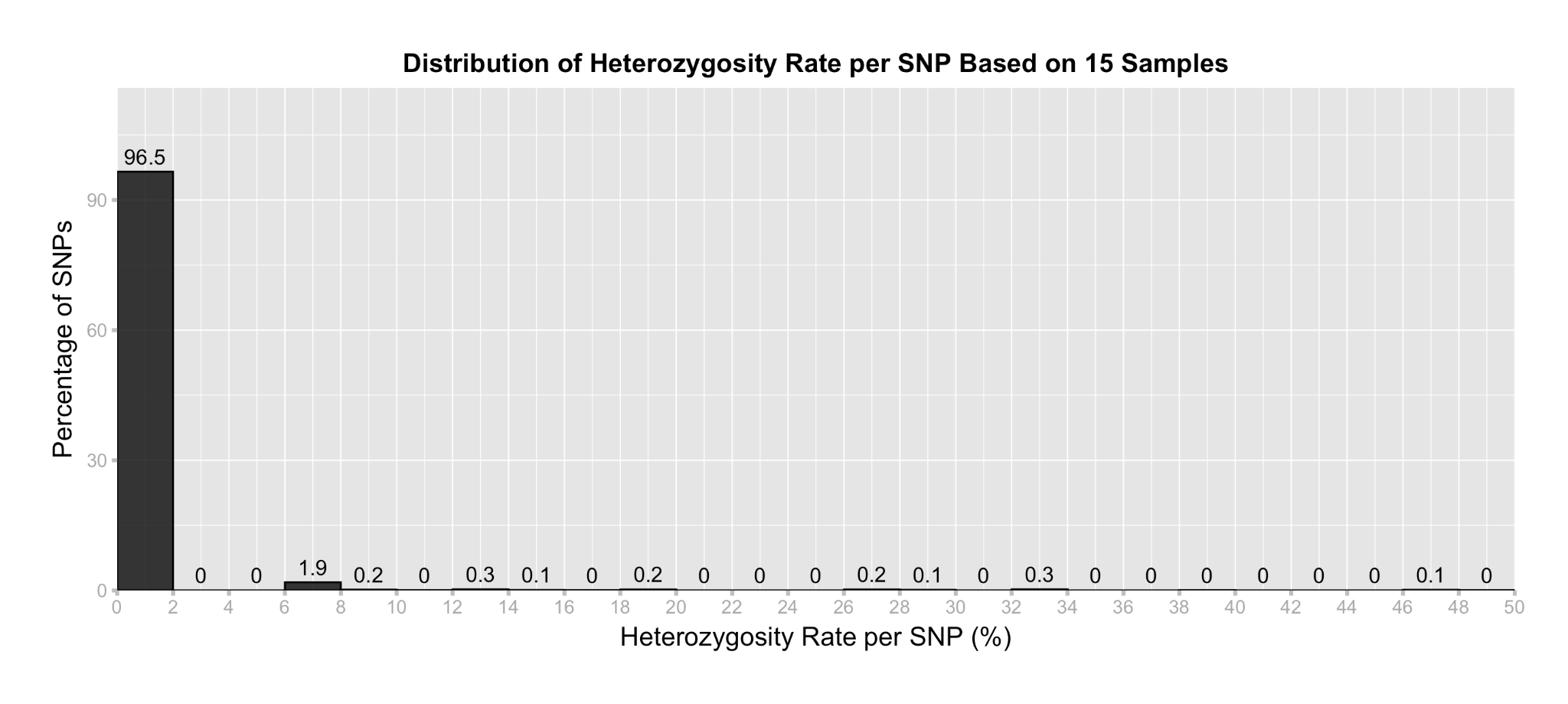

**Supplementary Figure 4.** Distribution of heterozygosity rate per SNP based on 2,421 SNPs in SCGA and 15 breeding accessions.
