## Supplementary Table for "Development and evaluation of a cost-effective, mid-density SNP array as a sorghum community genotyping resource"

**Supplementary Table 1.** Phenotypic trait-linked SNPs in the sorghum Agriplex array. The physical location of the phenotypic traits has not been independently confirmed (or supported) and should be considered as putative.

| **Chr** | **Pos (bp)** | **Trait** | **EIB’s Source** |
| --- | --- | --- | --- |
| 2 | 6045380 | Fertility restoration | S. Deshpande, Unpub. info. |
| 2 | 6843380 | Fertility restoration | S. Deshpande, Unpub. info. |
| 2 | 8715821 | Fertility restoration | S. Deshpande, Unpub. info. |
| 2 | 59000770 | Post-flowering Drought tolerance | S. Deshpande, Unpub. info. |
| 2 | 59821923 | Post-flowering Drought tolerance | S. Deshpande, Unpub. info. |
| 2 | 60098184 | Post-flowering Drought tolerance | S. Deshpande, Unpub. info. |
| 2 | 61811307 | Post-flowering Drought tolerance/Fertility restoration | S. Deshpande, Unpub. info. |
| 2 | 67306935 | Post-flowering Drought tolerance | S. Deshpande, Unpub. info. |
| 2 | 67710384 | Post-flowering Drought tolerance | S. Deshpande, Unpub. info. |
| 2 | 71419274 | Post-flowering Drought tolerance | S. Deshpande, Unpub. info. |
| 3 | 30310883 | Fertility restoration gene | S. Deshpande, Unpub. info. |
| 4 | 364279 | Fertility restoration gene | S. Deshpande, Unpub. info. |
| 5 | 838874 | Shoot fly resistance, Leaf Glossiness | S. Deshpande, Unpub. info. |
| 5 | 1608322 | Fertility restoration | S. Deshpande, Unpub. info. |
| 5 | 69794954 | Striga Resistance, StrR | Gobena et al. (2017) PNAS, 114(17): 4471-4476, https://doi.org/10.1073/pnas.1618965114 |
| 5 | 69847924 | Striga Resistance, StrR | Gobena et al. (2017) PNAS, 114(17): 4471-4476, https://doi.org/10.1073/pnas.1618965114 |
| 5 | 69851828 | Striga Resistance, StrR | Gobena et al. (2017) PNAS, 114(17): 4471-4476, https://doi.org/10.1073/pnas.1618965114 |
| 5 | 69852443 | Striga Resistance, StrR | Gobena et al. (2017) PNAS, 114(17): 4471-4476, https://doi.org/10.1073/pnas.1618965114 |
| 6 | 2682627 | Sugarcane Aphid Resistance, SCAR | T. Felderhoff, Unpub. info. |
| 6 | 2892438 | Sugarcane Aphid Resistance, SCAR | T. Felderhoff, Unpub. info. |
| 8 | 653850 | Partial Fertility restoration gene | S. Deshpande, Unpub. info. |
| 8 | 60934182 | Partial Fertility restoration gene | S. Deshpande, Unpub. info. |
| 9 | 4248350 | Fertility restoration gene | S. Deshpande, Unpub. info. |
| 9 | 46616891 | Fertility restoration gene | S. Deshpande, Unpub. info. |
| 10 | 58031253 | Shoot fly resistance, Trichome density | S. Deshpande, Unpub. info. |
| 10 | 60919657 | Shoot fly resistance, Trichome density | S. Deshpande, Unpub. info. |
